# Development of a continuous bioreactor to maintain stable nasal microbiomes from swab specimens and synthetic communities

**DOI:** 10.64898/2026.03.16.712028

**Authors:** Soyoung Ham, Marcelo Navarro-Diaz, Laura Camus, Timo N. Lucas, Paolo Stincone, Simon Heilbronner, Hannes Link, Daniel Petras, Daniel H. Huson, Largus T. Angenent

## Abstract

**Background:** The nasal microbiome is a collection of diverse microbial populations that inhabit the nose. *Staphylococcus aureus* is the most common opportunistic pathogen that colonizes the nasal mucosa, increasing the risk of invasive infections in immunocompromised and hospitalized patients. Clinicians usually prescribe antibiotics to decolonize the nasal cavities of at-risk patients from *S. aureus*. However, their broad antimicrobial activity can damage the resident nasal microbiome. Instead, naturally occurring compounds or resident bacteria in nasal microbiomes can effectively and safely exclude *S. aureus* from the nose. Cell culture and animal models have been used for nasal microbiome studies. However, their unstable microbiomes reduce the accuracy and reliability of the results. Recently, continuous bioreactors have been proposed as alternatives to these models.

**Results:** We designed and operated a continuous bioreactor system to maintain stable nasal microbiomes. Next, we inoculated the bioreactor with nasal-swab specimens that we had collected from healthy volunteers. We operated the bioreactors under varying conditions (*i.e.*, operating mode, dilution rate, temperature, pH, and medium composition), and determined the optimal conditions (continuous mode, 1 d^-1^, 30xlink:href=" pH 6.5, and synthetic nasal medium 3), resulting in stable microbiomes consisting of the main nasal bacterial species. The nasal microbiomes in the optimized bioreactors showed high reproducibility and resilience during a pH perturbation. Moreover, all microbiomes in the bioreactor, which were inoculated with six different nasal-swab specimens, maintained stable bacterial and metabolite compositions. In addition, we applied a synthetic microbial community (SynCom), which was derived from one of the volunteers, to demonstrate a *S. aureus* decolonization strategy. The bioreactor, inoculated with this SynCom, maintained a stable nasal microbiome for more than one month. Finally, different *S. aureus* strains that we inoculated in the SynCom showed distinct growth patterns within the otherwise stable community.

**Conclusions:** The continuous bioreactor enables the cultivation of stable nasal microbiomes for longer than one month by mimicking the environmental conditions of the human nose. The bioreactor is a valuable model for understanding the functions of the nasal microbiome and devising new decolonization strategies against *S. aureus*.

## 1. Background

The nasal microbiome is a consortium of microbial populations that reside in the human nose [1]. The human nose is an essential entry point connecting the external environment to the respiratory system and is, therefore, exposed to a wide variety of microbes, pollutants, and aeroallergens [2]. As a result, the nasal microbiome varies across individuals and is influenced by environmental factors (*e.g.*, birth mode, feeding method, siblings, antibiotic treatment, and smoking) [3]. The microbial composition and its resilience to change are important for human health [4], and *Staphylococcus*, *Corynebacterium*, *Propionibacterium*, and *Moraxella* are the major genera found in the human nose [5, 6]. This represents the co-existence of commensal and pathogenic bacteria through positive, negative, and neutral interactions [7, 8]. Pathogen dominance, such as with the relevant pathogen *Staphylococcus aureus*, has been linked to microbiota dysbiosis in chronic rhinosinusitis patients and to an increased risk of subsequent infection in healthy individuals [6, 9, 10].

*S. aureus* is a human opportunistic pathogen that asymptomatically colonizes different body sites, including the nose [11, 12] and the gut [13]. *S. aureus* is an efficient colonizer of nasal cavities due to its competitiveness regarding adhesion sites and nutrients, while resisting the activity of antimicrobial compounds and host immune responses [14]. These features may explain why *S. aureus* colonizes the nasal habitats of approximately 30% of the human population [15]. Albeit asymptomatic, *S. aureus* colonization poses a serious risk of invasive infections, particularly in immunocompromised and hospitalized patients [12]. Mupirocin is the most commonly used antibiotic for decolonizing the nose from *S. aureus* and preventing infections after surgery [16]. However, this antibiotic exhibits broad antibacterial activity and can, thereby, damage the nasal microbiome [17, 18]. Moreover, the presence of antibiotic-resistant *S. aureus* undermines the efficacy of mupirocin treatment [19]. To address these problems, naturally occurring compounds and resident bacteria in nasal microbiomes could be used to decolonize the nose from *S. aureus* as an alternative to antibiotic treatment.

*In-vitro* cell culture and animal models have been used in nasal microbiome studies [2, 20–23]. Culture models using plates and flasks are easy to handle but lack the complexity of the nasal environment. Notably, their closed system, with a fixed nutrient supply and inadequate pH control, promotes the dominance of a few microbial species, and thereby compromises the accuracy and reliability of the results. An alternative model could be animal models, including gnotobiotic animal models, but these exhibit distinct physiological properties compared to humans and are hindered by low reproducibility, ethical concerns, and cost limitations [24, 25]. Therefore, another model is needed, which more closely mimics the nasal environment and maintains stable microbiomes. With recent advances in microbiome science, continuous bioreactors are emerging as promising alternatives because they can reproduce steady-state conditions for environmental parameters, such as pH and oxygen concentration, from body sites [26, 27]. Moreover, bioreactors enable the stable cultivation of complex microbiomes throughout extended operating periods at higher cell densities than to *in-vitro* culture models [28, 29]. This enables us to explore ecological niches and resistance mechanisms against pathogen colonization. Nevertheless, most studies on using bioreactors to sustain microbiomes have focused on the human gut microbiome [25, 29–33]. To the best of our knowledge, a continuous bioreactor to cultivate nasal microbiomes has yet to be developed.

Lawson et al. proposed the design-build-test-learn cycle as a guideline for microbiome science, which can be utilized for bioreactor development [34]. This cycle involves rational design and building of microbiome models, testing their performance and functions, and learning from successes and failures to implement the models in subsequent cycles. In detail, two approaches (*i.e.*, top-down and bottom-up approaches) can be applied to bioreactor development [24]. In a top-down approach, human microbiomes (*e.g.*, nasal-swab specimens) are used to optimize bioreactors by evaluating various operating conditions. Conversely, a bottom-up approach entails the use of synthetic microbial communities (SynComs) to characterize the individual contributions to the microbiome functioning. The integration of the two approaches mitigates the risks of failure in the bioreactor development [27].

Our ultimate goal is to devise innovative strategies for *S. aureus* decolonization using a continuous bioreactor as a nasal model. Here, we divided the work into five experiments for two approaches. For the first four experiments, we developed a continuous bioreactor using a top-down approach by inoculating with nasal-swab specimens that were collected from healthy volunteers. For the first experiment (*i.e.*, optimization experiment), we optimized steady-state operating conditions by changing five conditions: **(1)** operating modes (batch *vs.* continuous); **(2)** dilution rate; **(3)** temperature; **(4)** pH; and **(5)** medium composition. At the optimized conditions, we then performed three further experiments: reproducibility, perturbation, and stabilization experiments with up to six nasal-swab specimens as inocula. For these three experiments, we analyzed optical densities, bacterial community compositions (*via* qPCR and sequencing of the 16S-ITS-23S rRNA gene operon), and metabolites (*via* non-targeted metabolomics). For the fifth experiment (*i.e.*, decolonization experiment), we switched to a bottom-up approach. We inoculated our bioreactor with a SynCom that we had assembled from nasal strains and that had been isolated from one of the volunteers. We then operated five bioreactors under our optimized operating conditions. We first maintained stable nasal microbiomes of the SynCom without *S. aureus*, and then added two different *S. aureus* strains to demonstrate *S. aureus* nasal decolonization strategies.

## 2. Methods

### 2.1. Preparation of nasal-swab specimens and a SynCom

We selected six healthy volunteers, regardless of gender or age (**Table S1**), who showed no signs of infection and had not received antibiotic treatment in the two weeks prior to sampling. We collected nasal-swab specimens from both nostrils using an ESwab collection and transport kit (BD Bioscience, San Jose, CA, USA). For each volunteer, we sampled nasal-swab specimens six times throughout two days, and then stored them at 4 until bioreactor inoculation. The clinical ethics committee of the University of Tübingen (No. 109/2009 BO2) approved the specimen-collection procedures. We designed a single SynCom from nine nasal strains that were isolated from Volunteer 1 and that were deposited in the LaCa collection (**Table S2**) [20]. The SynCom was prepared as described previously from precultures in rich medium of the different strains, as detailed in Table S2 [25]. Briefly, bacterial cells from precultures were washed and resuspended in a phosphate-buffered saline-glycerol solution to a final optical density at 600 nm (OD_600nm_) of 5. We then assembled the SynCom by combining the resuspended nasal strains in an equal ratio. Finally, we aliquoted and stored the completed SynCom at -80 until inoculation.

### 2.2. Bioreactor operation

We operated six 250-mL, glass bioreactors (DWK Life Sciences, Mainz, Germany) as axenic chemostats in a continuously-fed mode (**Fig. 1**). We maintained a fully stirred liquid within the bioreactors using a magnetic stirrer (2mag, München, Germany) at a 250-rpm setting to maintain homogeneous conditions and an oxygen flux into the liquid. We controlled the liquid temperature to 25–35 using a water-circulation bath (Huber, Offenburg, Germany) and a liquid jacket around the glass bioreactors. We controlled the pH to 5.5–7.5 by inserting a pH probe (Xylem, WA, USA), using a controller (Xylem), and pumping 0.5 M acid and 0.5 M base when needed (Cole-Parmer, Chicago, IL, USA). We maintained microaerobic conditions with a large air filter (Cytiva, Marlborough, MA, USA) and measured oxygen concentrations with a real-time fluorescing sensor (PreSens, Regensburg, Germany). We operated bioreactors in batch and continuous modes. For continuous mode, peristaltic pumps (Cole-Parmer) supplied synthetic nasal medium (SNM) at dilution rates 0.5–1.5 d^-1^ [14]. SNM is a synthetic minimal medium composed of the main metabolites from nasal habitats [14], which we varied in concentration for the bioreactor optimization study.

**Fig. 1.**
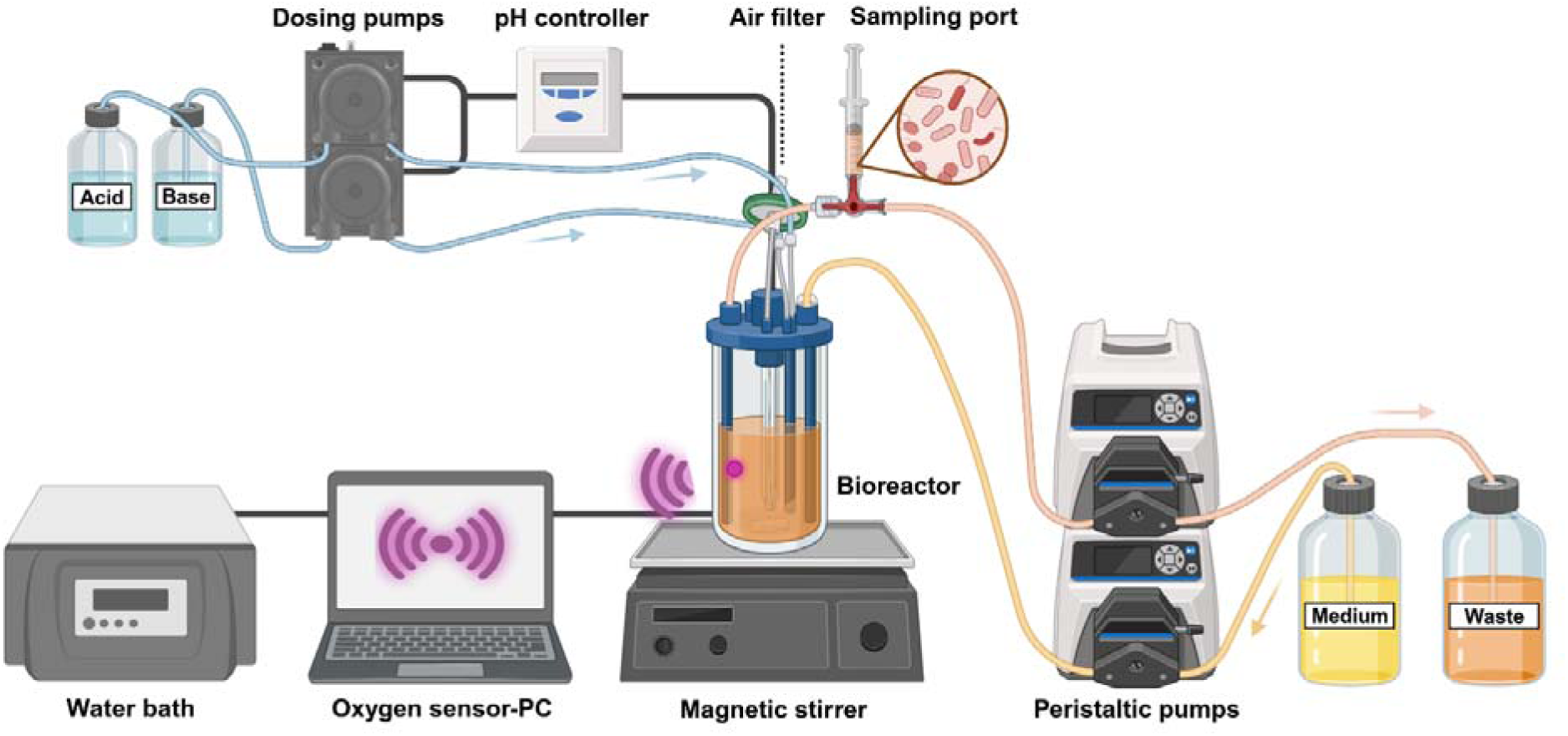
Schematic diagram of the continuous bioreactor setup to cultivate nasal microbiomes. Th graphic was created with BioRender.com.

We used either nasal-swab specimens or SynComs as inocula (**Table S3**). We inoculated 1 mL of fresh nasal-swab specimens or 2.5 mL of the SynCom (OD_600nm_=1) into the bioreactor on Day 0. For the decolonization experiment, we added two different *S. aureus* strains (*i.e.*, Sa C or Sa A) (Table S2) to the bioreactor on Day 20 (in duplicate) at a 1000-fold lower OD_600nm_ than the grown nasal microbiomes. We included one control bioreactor without *S. aureus* addition. We collected a 2-mL liquid sample from the nasal microbiomes daily through a sampling port (Braun, Hessen, Germany). We measured OD_600nm_ of the nasal microbiomes using a spectrophotometer (Thermo Scientific, Karlsruhe, Germany) and stored them at –20 for further experiments.

### 2.3. DNA extraction and qPCR

We centrifuged the liquid sample at 10,000 rpm for 10 min. After discarding the supernatants, we extracted DNA from the pellets according to the manufacturer’s instructions using the DNeasy PowerSoil Pro Kit (Qiagen, Hilden, Germany). We used a universal primer set (*i.e.*, 27-F and 342-R) to detect the 16S rRNA gene of total bacteria, and the *Sa442* gene-targeting primer set for *S. aureus* detection (**Table S4**) [35, 36]. We prepared a mixture containing 10 μL of SYBR Green PCR 2X master mix (Thermo Scientific), 1 μL of each forward and reverse primer (10 μM), 50 ng of DNA template, and sterilized distilled water to make a final volume of 20 μL. We aliquoted the mixture into wells in an optical 96-well fast clear reaction plate (Thermo Scientific). The thermal profile for the quantitative polymerase chain reaction (qPCR) was as follows: initial denaturation at 95 for 10 s, followed by 40 cycles of denaturation at 95 for 15 s, and annealing/extension at 60 for 60 s. We measured fluorescent signal intensities at the end of the annealing/extension step using a QuantStudio 3 (Thermo Scientific) and analyzed Ct values using Design & Analysis software (Thermo Scientific). We calculated the copy numbers of total bacteria and *S. aureus* from standard curves **(Figs. S1A–B)** [35]. We estimated qPCR efficiencies from the slopes of the standard curves that were reported in a previous study [37].

### 2.4. Amplicon sequencing

We targeted the 16S-ITS-23S rRNA gene operon (∼4,300 bp) for higher bacterial taxonomic resolution than the full-length 16S rRNA gene (∼1,500 bp) [38]. We mixed 50 ng of DNA template into 25 μL of LongAmp Taq 2X master mix (New England Biolabs, Ipswich, MA, USA) with 0.4 μM primers: 16S-27F-GGTGCTG-ONT (TTAACCTAGRGTTYGATYHTGGCTCAG) and 23S-2241R-GGTGCTG-ONT (TTAACCTACCRCCCCAGTHRAACT) [38]. We amplified the bacterial rRNA operon using optimized thermal conditions on the Mastercycler Pro (Eppendorf, Hamburg, Germany): an initial denaturation of 1 min at 94xlink:href=" followed by 30 cycles of 30 s at 94xlink:href=" 30 s at 52xlink:href=" and 1 min at 65xlink:href=" with a final step of 10 min at 65 . We cleaned PCR amplicons with AMPure XP (Beckman Coulter, Brea, CA, USA) and confirmed their size by agarose gel electrophoresis. We quantified the cleaned amplicons using a Qubit™ 1X dsDNA high-sensitivity kit (Invitrogen, Carlsbad, CA, USA). We prepared a sequencing library according to the standard procedure of a rapid barcoding kit (SQK-RBK114, Oxford Nanopore Technologies, Oxford, UK). We loaded the library onto a flow cell (R10.4.1, Oxford Nanopore Technologies) and sequenced for 12 h using a MinIon nanopore sequencer (Mk1B, Oxford Nanopore Technologies).

We used MMonitor (https://mmonitor.org) to analyze sequencing data [39]. Briefly, we performed taxonomic profiling using Emu v3.4.5 and the standard database that is included with the software, which combines NCBI RefSeq 16S rRNA gene sequences (downloaded January 2023) and rrnDB v5.8 sequences [40]. We performed principal coordinates analysis (PCA) by the GraphPad Prism v9.5.0 based on parallel analysis.

### 2.5. Non-targeted metabolomics

To extract metabolites from the liquid samples, we suspended pellets in 1 mL of 80% methanol (Thermo Scientific). We sonicated the suspensions for five cycles (60 s of operation with 40% amplitude, followed by 10 s of pause) using a sonicator (Thermo Scientific). We transferred supernatants obtained from centrifugation (10,000 rpm for 10 min) to a pre-weighed glass vial. We evaporated methanol in the vial using a Plus vacuum concentrator (Eppendorf) and weighed the pellets before and after evaporation. We resuspended the evaporated samples with 80% methanol to achieve a 5-mg µL^-1^ concentration.

We conducted non-targeted metabolomics using an UHPLC connected to a Q-Exactive HF mass spectrometer and a C18 core-shell column (Kinetax, 50 × 2.1 mm, 1.7 µm particle size, 100 Å pore size) (Phenomenex, Torrance, USA). We used two solvents: **(1)** water (Thermo Scientific) + 0.1% formic acid; and **(2)** acetonitrile (Thermo Scientific) + 0.1% formic acid. After sample injection, we performed a linear gradient method for 10 min at a flow rate of 500 µL min^-1^. We measured the samples in positive mode using heated electrospray ionization (HESI) with sheath gas at 50 AU, auxiliary gas at 12 AU^1^, and sweep gas at 1 AU. We set the spray voltage to 3.5 kV and the inlet capillary temperature to 250xlink:href=" while we set the S-lens RF level to 50 V and the auxiliary gas heater temperature to 400 . A full mass spectrometry (MS) 1 survey scan acquisition range was 120–1,500 m/z, with 60,000 resolution, 1E6 of automatic gain control (AGC), and a maximum injection time of 100 ms. We acquired the MS2 spectra with data-dependent acquisition mode by modifying a reported method [41]. We set the MS2 resolution to 15,000, the AGC target to 5E5, and the maximum injection time to 50 ms. We subjected the five most abundant precursor ions to MS/MS fragmentation. We used a precursor ion isolation window at 1 m/z and performed fragmentation with a normalized collision energy applied in stepped gradients of 25, 35, and 45. We scanned MS2 at the apex of peaks for 2–15 s with 5 s of dynamic exclusion. We selected the 100 statistically most significant metabolites using a random forest approach implemented in the FBMN-STATS software [42]. We scaled the intensities of the selected metabolites’ features using min-max normalisation to the range -1 to 1 [43]. For a heatmap, we used the ComplexHeatmap package for R v4.4.0 (RStudio v2024.12.1) [43]. We performed GraphPad Prism v9.5.0 for graphical and statistical analysis.

## 3. Results

### 3.1. Optimization of the bioreactor operating conditions to cultivate nasal microbiomes – optimization experiment

We collected nasal swab specimens from six volunteers without apparent disease symptoms or antibiotic treatment (**Table S1**). The nasal-swab specimens from our volunteers exhibited diverse bacterial community compositions (**Figs. S2A–B**). We classified the six nasal-swab specimens into three groups based on the most abundant genera: nasal-swab specimens from Volunteers 1 and 2 with *Staphylococcus* spp., those from Volunteers 3 and 4 with *Corynebacterium* spp., and those from Volunteers 5 and 6 with *Moraxella* spp. (**Fig. S2A, Table S1**) [44]. Volunteers 1 and 2 were dominated by *S. aureus* with a relative abundance of 29–62%, while relative *S. aureu* abundance was below 5% in Volunteers 3–6 (**Fig. S2B, Table S1**). We decided to inoculate a nasal-swab specimen from Volunteer 1 into the bioreactor for the optimization experiment because its bacterial composition showed a relatively higher abundance of *S. aureus* than that of other volunteers. One of our aims was to determine whether *S. aureus* would dominate the microbiome in bioreactors under batch- or continuous-culture conditions. The nasal-swab specimen from Volunteer 1 also comprised common nasal genera (*e.g.*, *Staphylococcus*, *Corynebacterium*, and *Cutibacterium*) **(Figs. S2A–B)** [5, 6].

During the optimization experiment, we varied operating conditions. First, we compared bioreactors that were operated in batch and continuous modes **(Figs. 2A–B)**. After Day 1 of the operating period in batch mode, relative *S. aureus* abundance in nasal microbiomes was nearly 100% and did not change throughout the period (**Fig. 2A**). In contrast, *Staphylococcus epidermidis* abundance increased in the continuous mode from Day 3 (**Fig. 2B**). The bacterial compositions became stable after Day 8 of the operating period, with approximately 82% of *S. epidermidis*, 15% of *S. aureus,* and 3% of *Staphylococcus capitis* (**Fig. 2B**). The OD_600nm_ in the batch bioreactor slightly decreased during time, while it remained constant in the continuous bioreactor within the range of 0.41–0.47 from Day 2 (**Figs. 2C, S3A**).

**Fig. 2.**
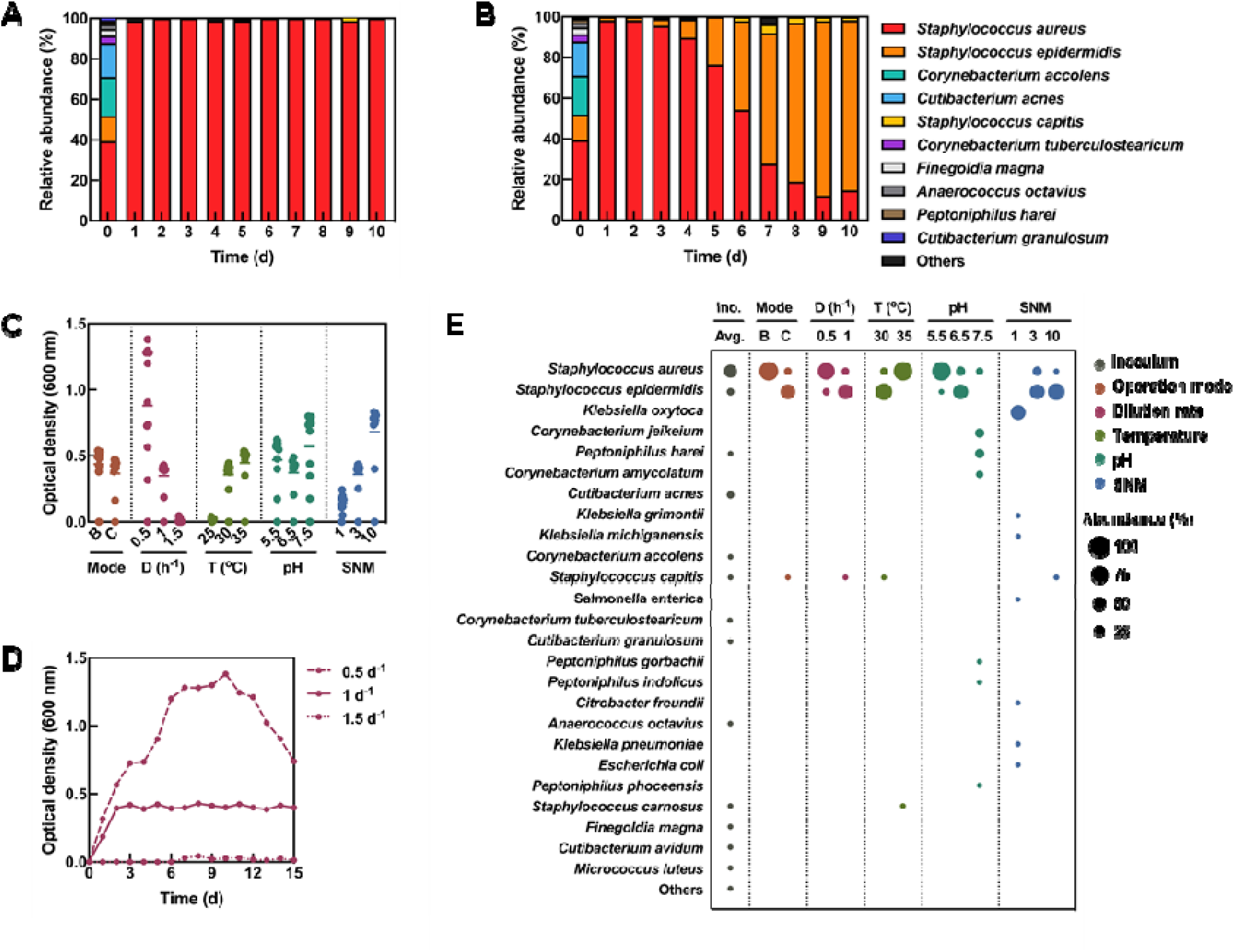
Optimization experiment for the bioreactors under various operating conditions. **A–B.** Comparison of bacterial community compositions of nasal microbiomes in batch (**A**) and continuous (**B**) modes. Bacterial composition of the nasal-swab specimen is shown on Day 0 as an initial inoculum. The order and color of species for panels **A** and **B** are the same and are based on the combined average relative abundance throughout the operating period for both bioreactors. The most abundant species are placed from bottom to top of the bar, but their names are listed from top to bottom in the legend. The remaining species are grouped under ‘others’ when not in the top ten. **C.** Optical densities of nasal microbiomes throughout the operating period under various conditions (operating mode, dilution rate, temperature, pH, and medium composition). Solid lines indicate the mean value. **D.** Optical densities of nasal microbiomes throughout the operating period at different dilution rates. **E.** Bubble plot showing bacterial community compositions of nasal-swab specimen (Ino.) and nasal microbiomes at the end of the operating period. Ino.: inoculum, B: batch mode, C: continuous mode, D: dilution rate, T: temperature, SNM: synthetic nasal medium.

Second, we tested dilution rates of 0.5, 1, and 1.5 d^-1^ with associated flow rates of 0.09, 0.18, and 0.26 mL min^-1^, respectively, for 250-mL bioreactors (**Figs. 2C–D, S3B**). The average OD_600nm_ was below 0.04 at the highest dilution rate of 1.5 d^-1^, because the microbiomes were washed out of the bioreactors (**Figs. 2C–D**) [25]. The average OD_600nm_ was the highest at our lowest dilution rate of 0.5 d^-1^ (**Fig. 2D**). Third, we changed the temperatures and observed that the nasal microbiomes did not grow at 25xlink:href=" and that the average OD_600nm_ at 30 and 35 were 0.36 and 0.45, respectively (**Figs. 2C, S3C**). Fourth, changes in pH also led to differences in average OD_600nm_, ranging from 0.37–0.57 at a pH of 7.5 to the highest OD (**Figs. 2C, S3D**). Fifth, to investigate the effects of medium composition, we utilized SNM 1, 3, and 10 with different concentrations of glucose, amino acids, and organic acids (**Figs. 2C, S3E**) [14]. As expected, OD_600nm_ increased with increasing nutrient concentrations in the SNM (**Figs. 2C, S3E**). Collectively, the average OD_600nm_ varied across conditions but remained stable after Day 6 of the operating period, except at batch mode and 0.5 d^-1^ (**Figs. S3A–E**).

After sampling the nasal microbiomes at the end of the operating period for these five operating conditions, we stored them at -20 and analyzed their bacterial community compositions using amplicon sequencing (**Figs. 2E, S4A–E**). Overall, the most abundant species were *S. aureus* and *S. epidermidis* (**Fig. 2E**). In particular, we found *S. aureus* to have reached a higher than 97% relative abundance, which is an unfavorable feature, under the following conditions: batch mode, 35xlink:href=" and pH 5.5 **(Figs. S4A, C–D)**. However, *S. aureus* did not dominate under other conditions, and it did not grow in SNM 1 (**Figs. S4A–E**). qPCR analysis with copy numbers of *Sa442* genes supported the *S. aureus* abundance patterns in the bioreactor (**Figs. S5A–E**) [36]. Instead, other nasal genera (*e.g.*, *Klebsiella*, *Corynebacterium*, *Peptoniphilus,* and *Cutibacterium*) accounted for 34–93% of the nasal microbiomes depending on operating conditions (**Fig. 2E**). Therefore, the following conditions, namely, a continuous mode, a dilution rate of 1 d^-1^, a temperature of 30xlink:href=" a pH of 6.5, and a medium concentration with SNM 3, were considered to be optimal. This is because they simultaneously satisfied three requirements: **(1)** the nasal microbiomes grew steadily within the bioreactor; **(2)** the community composition of the nasal microbiomes exhibited diversity consisting of main nasal bacteria; and **(3)** the stable communities included but were not dominated by *S. aureus*.

### 3.2. Reproducibility of nasal microbiomes in the continuous bioreactor – reproducibility experiment

Using the five optimized conditions, we operated six bioreactor replicates, with inoculation as the only difference. To test the reproducibility of our setup, we inoculated six different nasal-swab specimens (*i.e.*, nasal-swab specimens 1–6) from Volunteer 1, which were collected repeatedly throughout two days (see Methods). Specifically, we used the terminology *Nasal microbiome 1***–***6* to refer to the nasal microbiomes after inoculation with nasal-swab specimens 1**–**6, respectively (**Figs. 3A–E**). These nasal-swab specimens 1–6 exhibited only minor differences in species composition and relative abundance through 16S-ITS-23S rRNA gene sequencing (**Fig. 3A**). At the end of the operating period, Nasal microbiomes 1–6 were almost identical and were dominated by *Staphylococcus* spp. (**Fig. 3B**). When comparing bacterial compositions between the Nasal microbiomes 1 *and* 6 from the bioreactors, for example, both the Pearson correlation coefficient (r) and the coefficient of determination (R^2^) were higher than between the nasal-swab specimens 1 *and* 6 **(Figs. S6A–B)**. This revealed a stronger linear correlation and better model fit for the nasal microbiomes grown in the bioreactor than for the nasal-swab specimens, suggesting a more similar community structure. We observed small fluctuations in the Nasal microbiomes 1 *and* 6 during the adaptation phase (Days 2–6) (**Figs. S6C–D**). The communities eventually reached the stable microbiomes (> Day 8) with similar relative abundances: 25–30% of *S. aureus*, 68–74% of *S. epidermidis*, and 1–3% of *S. capitis* for both bioreactors (**Figs. S6C–D**). Furthermore, the nasal microbiomes of all six bioreactors exhibited similar patterns in OD_600nm_ and copy numbers of total bacteria and *S. aureus* throughout the operating period (**Figs. 3C–E**). Thus, our bioreactors demonstrated high reproducibility under optimized operating conditions.

**Fig. 3.**
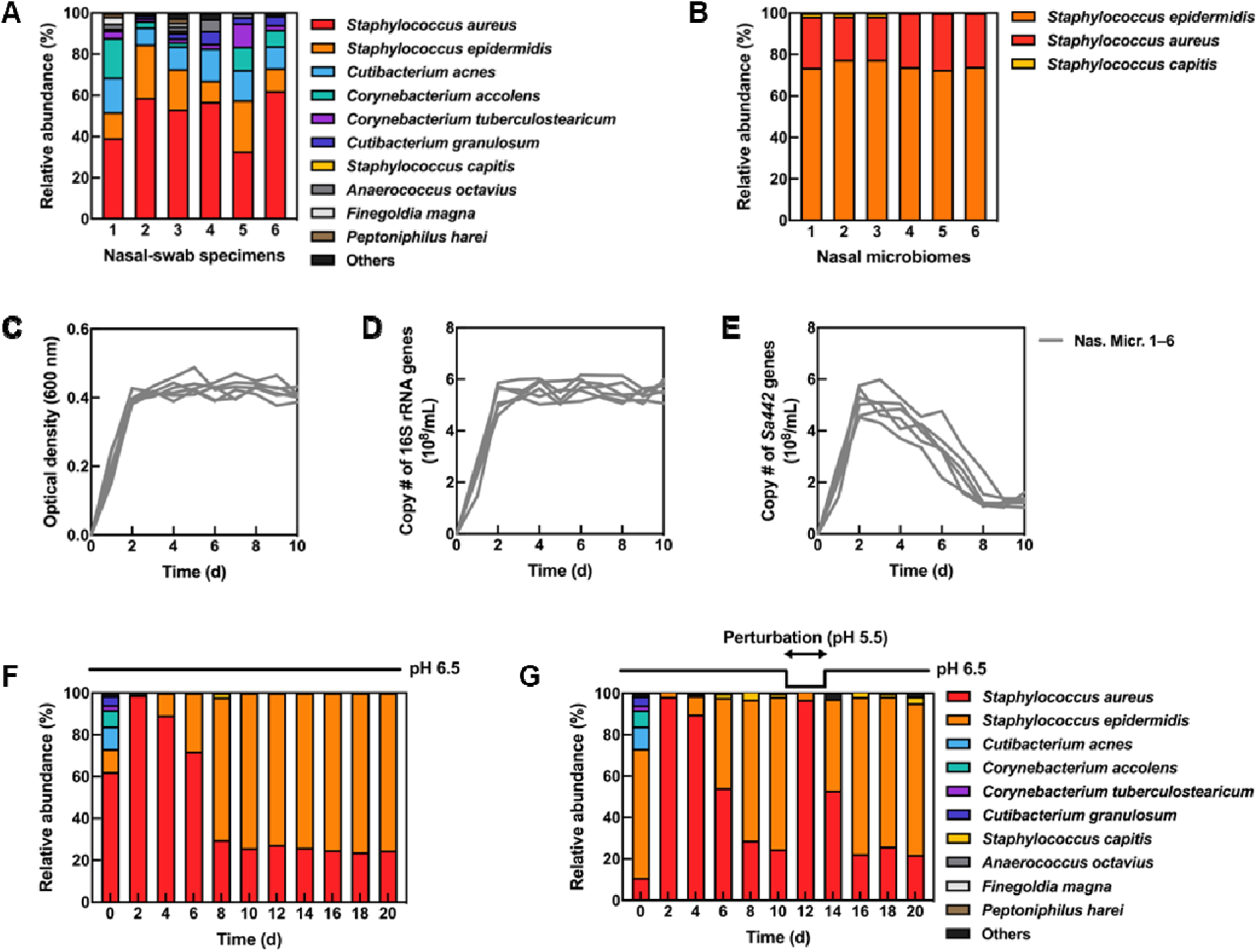
Reproducibility and perturbation experiments of the continuous bioreactor. **A–B.** Bacterial community compositions of nasal-swab specimens 1–6 that were derived from Volunteer 1 (**A**) and Nasal microbiomes 1–6 in the bioreactor at the end of the operating period (**B**) for the reproducibility experiment. The order of species is based on the average relative abundance for the six nasal-swab specimens **(A)** and the six nasal microbiomes (**B**), respectively. The most abundant species are placed from bottom to top of the bar, but their names are listed from top to bottom in the legend. However, the color for each species is the same between the panels **A–B**. **C–E.** Optical densities (**C**), copy numbers of total bacteria (**D**), and copy numbers of *S. aureus* (**E**) of nasal microbiomes throughout the operating period for the reproducibility experiment. **F–G.** Changes in bacterial community compositions of nasal microbiomes in two parallel bioreactors inoculated with nasal-swap specimen 6 following a pH perturbation (perturbation experiment). One bioreactor was not perturbed (**F**), while the other bioreactor was subject to a pH perturbation (**G**). Bacterial composition of the nasal-swab specimen is shown on Day 0 as an initial inoculum. The order and color of species for panels **F–G** are the same and are based on the combined average relative abundance throughout the operating time for both bioreactors. The most abundant species are placed from bottom to top of the bar, but their names are listed from top to bottom in the legend. The remaining genera or species are grouped under ‘others’ when not in the top ten. Nas. Micr.: Nasal microbiome.

### 3.3. Resistance and resilience of nasal microbiomes in the continuous bioreactor – perturbation experiment

pH levels can fluctuate frequently in nasal habitats [45, 46], and such fluctuations strongly affect the stability and resilience of microbial communities under stressful conditions [47, 48]. Therefore, we investigated the dynamics of bacterial community composition by temporarily perturbing the nasal microbiome pH (**Figs. 3F–G**). We inoculated the nasal-swab specimen 6 into two parallel bioreactors that were operated at a pH of 6.5 (**Figs. 3F–G**). We maintained the pH at 6.5 in one bioreactor (**Fig. 3F**), while perturbing it in the other bioreactor (**Fig. 3G**). To do this, we let the community stabilize until Day 10 at a pH of 6.5 after which we adjusted the pH to 5.5 for two days and then restored it to its original value (**Fig. 3G**). At a lower pH, *S. aureus* began to dominate, consistent with our previous results from the optimization experiment (**Fig. S4D**). The changes in the composition following our pH perturbation suggest that the nasal microbiome has limited resistance to the pH fluctuation (**Fig. 3G**). However, the nasal microbiomes rapidly recovered to their original state within two days (**Fig. 3G**). This highlights the strong resilience of the specific nasal microbiomes in our continuous bioreactor against pH perturbations. However, a universal conclusion would need additional experiments with other nasal microbiomes.

### 3.4. Stable nasal microbiomes in the continuous bioreactor – stability experiment

To investigate the dynamics of other nasal microbiomes in our bioreactor system, we inoculated the nasal-swab specimens from six volunteers (Volunteers 1**–**6) into separate bioreactors and operated them at the optimized conditions for 20 days, which, for this stability experiment, extended the operating period for 10 days compared to the optimization experiment (**Table S3**). The six nasal microbiomes in the bioreactors exhibited different trends in OD_600nm_ and in the copy numbers of total bacteria and *S. aureus*, but they all reached a stable community structure before the end of the operating period (**Figs. 4A–F, S7A–C**). Specifically for this stability experiment, we used the terminology *Nasal microbiome A***–***F* to refer to the nasal microbiomes after inoculation with nasal-swab specimens from Volunteers 1**–**6, respectively (**Figs. 4A–F**).

**Fig. 4.**
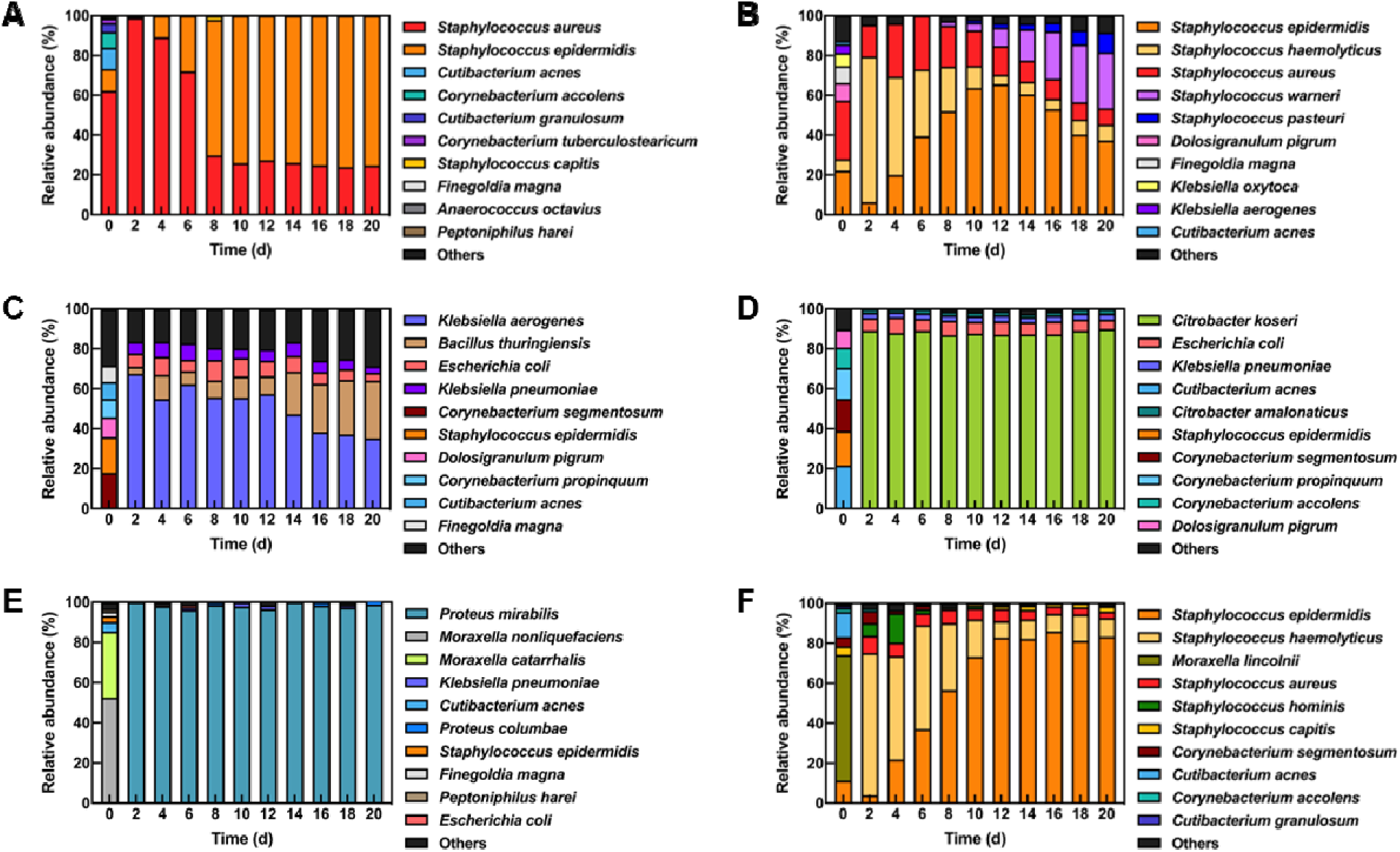
The stability of bacterial community experiment of the continuous bioreactor. Bacterial community compositions of nasal microbiomes in the bioreactor inoculated with six nasal-swab specimens (Nasal microbiomes A–F) for the stability experiment. **A–F.** Relative abundances of the top ten species in the Nasal microbiome A (**A**), Nasal microbiome B (**B**), Nasal microbiome C (**C**), Nasal microbiome D (**D**), Nasal microbiome E (**E**), and Nasal microbiome F (**F**) throughout the operating period. Bacterial composition of the nasal-swab specimens is shown on Day 0 as an initial inoculum. The order of species differs for each panel and is based on the combined average relative abundance throughout the operating period for each bioreactor. The most abundant species are placed from bottom to top of the bar, but their names are listed from top to bottom in the legend. However, the color for each species is the same between the panel **A–F**. The remaining species are grouped under ‘others’ when not in the top ten.

The majority of the Nasal microbiomes A and B consisted of *Staphylococcus* spp., reaching at least a 94% abundance from Day 2 of the operating period (**Figs. S8A–B**). For the Nasal microbiome A, *S. aureus* was the most abundant species until Day 6 (**Fig. 4A**). Subsequently, *S. epidermidis* began to increase to reach 76% on Day 8 after which we observed constant abundances of *S. aureus* and *S. epidermidis* (**Fig. 4A**). In contrast, we observed additional species of *Staphylococcus* (*e.g.*, *Staphylococcus haemolyticus*, *Staphylococcus warneri*, and *Staphylococcus pasteuri*) at high abundances in the Nasal microbiomes B and throughout the operating period (**Fig. 4B**). The relative abundances of *S. haemolyticus*, *S. warneri*, and *S. pasteuri* increased steadily from Day 12 of the operating period, whereas those of *S. aureus* and *S. epidermidis* decreased (**Fig. 4B**). At the end of the operating period for the Nasal microbiome B, all 5 species of *Staphylococcus* were present at a moderate (10%, *S. pasteuri*) to high (37%, *S. epidermidis*) relative abundance, showing a considerably higher diversity than for the Nasal microbiome A.

Nasal-swab specimens from Volunteers 3 and 4 were composed of *Corynebacterium* spp. and other genera (**Fig. S2A**). Whereas, for the Nasal microbiome C, *Klebsiella* spp. and *Bacillus* spp. constituted the most abundant genera. After stabilization of the Nasal microbiome C on Day 16, *Klebsiella* spp. and *Bacillus* spp. reached a composition of 39–44% and 38–46%, respectively (**Figs. 4C, S8C**). We found that the Nasal microbiome D composition remained stable from Day 2 onward during the operating period. The average relative abundances from Days 2–20 were 90–94% for *Citrobacter* spp., 4–8% for *Escherichia* spp., and 2–3% for *Klebsiella* spp. (**Figs. 4D, S8D**).

Despite the high relative abundance of *Moraxella* spp. (63–89%) for nasal-swab specimens of Volunteers 5 and 6, we could not cultivate them in the bioreactor (**Figs. S2A, S8E–F**). Instead, *Proteus* spp. and *Staphylococcus* spp. comprised most of the Nasal microbiome E and F (**Figs. S8E–F**). We detected *Proteus mirabilis* at a relative abundance of almost 100% in the Nasal microbiome E throughout the operating period (**Fig. 4E**). Relative *S. epidermidis* abundance increased from 11% in the inoculum to 81–83% in the Nasal microbiome F on Day 12 after which it remained stable. The rest of this Nasal microbiome F consisted mostly of *S. haemolyticus* (3–13%) and *S. aureus* (4–6%) (**Fig. 4F**). For the Nasal microbiomes C and E, we observed that species that were relatively low in abundance in the nasal-swab specimens became abundant (**Figs. 4C–E**).

In addition, we investigated metabolic profiles within Nasal microbiomes A**–**F after stabilization in the bioreactor, which occurred by Day 16 at the latest for Nasal microbiome B. We sampled and analyzed the nasal microbiomes on Days 16, 18, and 20 of the operating periods (**Figs. 4A–F**). We identified 3,086 metabolites using non-targeted metabolomics. We selected the 100 statistically most significant annotated metabolites and displayed their distributions on a heatmap (**Fig. 5A**). The composition of 100 metabolites for each of the six Nasal microbiomes A**–**F were unique, even though the metabolites of the Nasal microbiomes A and B were not completely separated (**Fig. 5A**). In addition, each nasal microbiome showed similar metabolic distributions on Days 16, 18, and 20 (**Fig. 5A**), suggesting that metabolite profiles remained stable toward the end of the operating period, mirroring the stability of the nasal microbiome composition for the stability experiment (**Figs. 4A–F**).

**Fig. 5.**
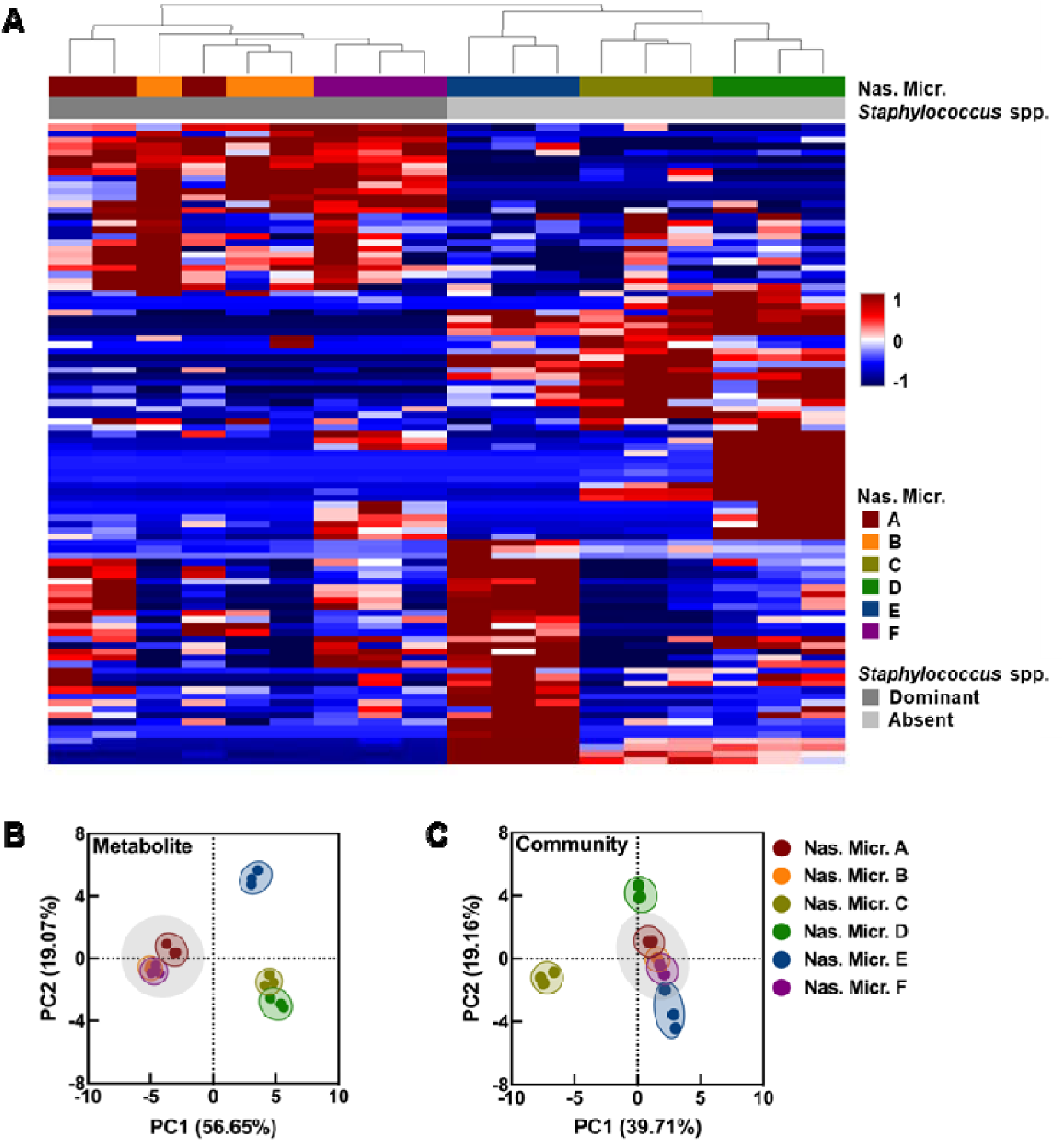
The stability of metabolic experiment of the continuous bioreactor. Metabolomic profile of nasal microbiomes in the bioreactors inoculated with six nasal-swab specimens (Nasal microbiomes A–F) for the stability experiment. Stable Nasal microbiomes A–F on Days 16, 18, and 20 were used for the metabolic profiles. **A.** Heatmap and the hierarchical clustering of statistically significant selected metabolites (100 in total). The intensity of each metabolite i scaled between -1 (blue) to 1 (red). The columns above the heatmap indicate the distribution of Nasal microbiomes A–F or the dominance of *Staphylococcus* spp. (Staphylococcal-dominant group *vs*. Staphylococcal-absent group). **B–C.** The comparison of PCA plots for the compositions of metabolites (**B**) and bacterial communities (**C**). Each dot represents one nasal microbiome, and its color indicates the variation of Nasal microbiomes A–F. A Staphylococcal-dominant group (Nasal microbiomes A, B, and F) is clustered with a gray background. Nas. Micr.: Nasal microbiome.

The hierarchical clustering analysis of these metabolic profiles revealed two distinct groups: Nasal microbiomes A, B, and F *vs.* Nasal microbiomes C**–**E (**Fig. 5A**). This binary, two-group structure reflected the dominance of *Staphylococcus* spp. (*i.e*., Staphylococcal-dominant group) in the stable Nasal microbiomes A, B, and F (**Figs. 4A, B, F**) *vs*. the absence of *Staphylococcus* spp. (*i.e*., Staphylococcal-absent group) in the stable Nasal microbiomes C**–**E (**Figs. 4C–E**). Thus, our observations suggest a close connection between the metabolite and species compositions of bacterial communities (**Figs. 4A–F, 5A**). A PCA plot of the metabolic profiles clearly supported a binary clustering into the Staphylococcal-dominant and Staphylococcal-absent groups (**Fig. 5B**). A PCA plot of the bacterial community composition did not show such clear binary clustering, even though clustering was observed: **(1)** the Staphylococcal-dominant group was tightly clustered due to the domination of the closely-related *Staphylococcus* spp.; and **(2)** the Staphylococcal-absent group was composed of bacterial communities that were more diverse, reflecting a larger distance between the communities (**Figs. 4A–F, 5C**).

### 3.5. *S. aureus* decolonization strategies using the continuous bioreactor – decolonization experiment

We demonstrated that our bioreactor system supports the stable cultivation of nasal microbial communities, with varying *S. aureus* abundances. Here, we investigated whether *S. aureus* proliferation within these communities is strain- and/or community-dependent, with the aim of establishing a robust model to evaluate decolonization strategies. For this decolonization experiment, we inoculated a SynCom rather than a nasal swab specimen, enabling precise control over the community members. During the operating run, we also added two different *S. aureus* strains to demonstrate that we can prevent *S. aureus* dominance in nasal microbiomes, and thus identify a possible decolonization strategy. We used the previously established SynCom C from Camus et al. [20], whose strains were originally isolated from Volunteer 1 in our study. This SynCom comprised nine strains, which were mixed in equal proportions, and excluded the native *S. aureus* strain from this Volunteer 1 (**Fig. 6A, Table S2**). We inoculated five bioreactors with SynCom C on Day 0 and then operated them until Day 20 to maintain stable nasal microbiomes (**Fig. 6A**). Following stabilization, we added to duplicate bioreactors two *S. aureus* strains that were isolated from the nasal-swab specimens of two volunteers: Sa C from Volunteer C (our Volunteer 1) and Sa A from Volunteer A (who was not used in our study) (**Fig. 6A, Table S2**) [20]. We operated the five bioreactors in three conditions: **(1)** one bioreactor with SynCom C as an inoculum alone as a negative control; **(2)** two bioreactors for SynCom C+Sa C; and **(3)** two bioreactors for SynCom C+Sa A (**Figs. 6B–F**).

**Fig. 6.**
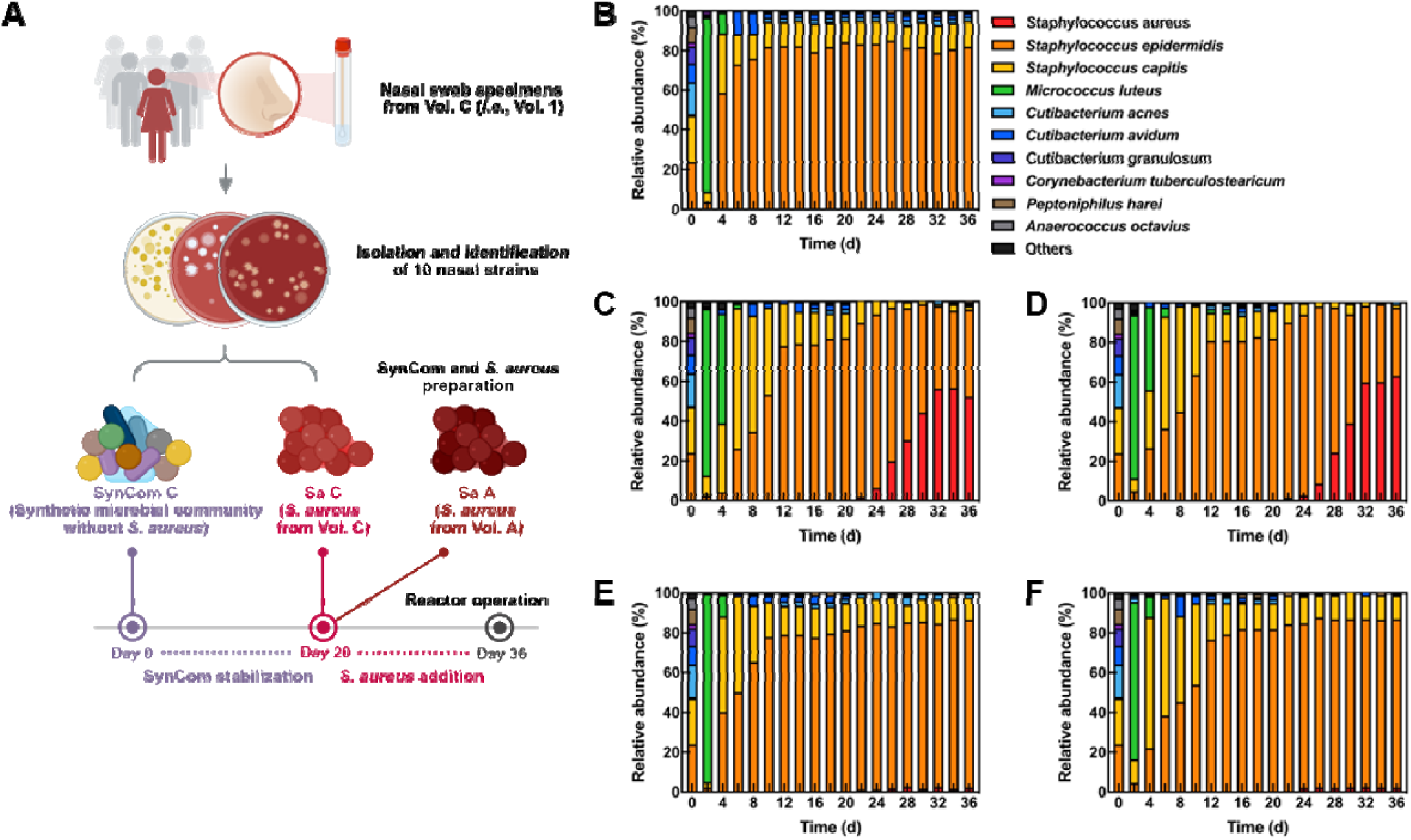
*S. aureus* decolonization experiment of the continuous bioreactor. Nasal microbiome changes in the bioreactor inoculated with SynCom and different *S. aureus* strains for the decolonization experiment. **A.** Schematic overview of the methodology for bioreactor operation. SynCom C was inoculated into the bioreactor on Day 0, and two different *S. aureus* strains (Sa C or Sa A) were added on Day 20. SynCom C was derived from Volunteer C (*i.e.*, Volunteer 1). Sa C was isolated from a nasal-swab specimen from Volunteer C, whereas Sa A was from Volunteer A. The graphic was created with BioRender.com. **B–F.** Relative abundances of nasal microbiomes in the bioreactor inoculated with SynCom C and two different *S. aureus* strains (Sa C or Sa A). Nasal microbiomes in the bioreactor inoculated with SynCom C alone (**B**). Nasal microbiomes in the bioreactors inoculated with SynCom C+Sa C (**C–D**) or SynCom C+Sa A (**E–F**). Duplicate bioreactors were operated to confirm the reproducibility of the results with *S. aureus* addition. The bacterial composition of SynCom C is shown on Day 0 as the initial inoculum. The order and color of species for panels **B–F** are the same. *S. aureus* is placed at the bottom of the bar, but its name is listed at the top in the legend. The rest of the species are based on the combined average relative abundance throughout the operating period for all bioreactors. The most abundant species are placed from bottom to top of the bar, but their names are listed from top to bottom in the legend. The remaining species are grouped under ‘others’ when not in the SynCom C.

OD_600nm_ and copy numbers, which represent the concentration of total bacteria in the nasal microbiomes, were similar in all bioreactors and reached a stable level after Day 8 of the operating period (**Figs. S9A–B**). The composition of the nasal microbiomes, which were inoculated with SynCom C alone (*i.e.*, negative control), became stable from Day 10 of the operating period with approximately 80% of *S. epidermidis*, 13% *S. capitis*, and 5% *Cutibacterium* spp. (**Fig. 6B**). As anticipated, no growth of *S. aureus* was observed due to the absence of this species as a negative control (**Fig. 6B)**.

We observed a dramatic difference in *S. aureus* growth based on whether *S. aureus* strains *was* (Sa C) or *was not* (Sa A) from the same volunteer as the other nasal strains in the SynCom C. Sa C started to dominate the nasal microbiome from Day 24, occupying 52–63% of stable nasal microbiomes after Day 32 of the operating period (**Figs. 6C–D, S9C**). Overall, the diversity of the nasal microbiomes decreased in response to the growth of Sa C, with a loss of *Cutibacterium* spp. (**Figs. 6C–D**). At the end of the operating period, the most abundant species were *S. aureus* and *S. epidermidis*, as observed in the nasal-swab specimen from the same volunteer (*i.e.*, Volunteer C) (**Figs. 4A, 6C–D**). In contrast, Sa A rarely appeared throughout the operating period (**Figs. 6E–F, S9C**). These results were confirmed with our bioreactor duplicates, although small fluctuations were observed only during the adaptation phase (Days 2–10) (**Figs. 6C–F**). Thus, replicated experiments showed that one decolonization strategy could be to alter the microbial community with commensal strains that had not been found with a specific *S. aureus* strain in volunteers, because Sa A did not proliferate in SynCom C, while Sa C did (**Figs. 6E–F** *vs*. **Figs. 6C–D**).

## 4. Discussion

### 4.1. Our continuous bioreactor cultivates stable nasal microbiomes for an extended operating period

Each human body site has unique environmental conditions (*e.g.*, pH, temperature, oxygen concentration, and nutrient availability), which contribute to distinct microbial populations [49]. The human nose is slightly acidic and has a relatively low temperature, similar to that of human skin, but with reduced oxygen concentrations [49]. Based on these conditions, we operated bioreactors with pH levels ranging from 5.5 to 7.5 and temperatures ranging from 25 to 35xlink:href=" while maintaining microaerobic conditions (**Figs. 1, 2A–E**) [25, 46, 50]. As a nutrient medium, we used SNM, which contains the main metabolites of nasal secretions and mimics the nasal environment [14].

Most importantly, we continuously cultivated nasal microbiomes in the bioreactor for an extended operating period of up to 36 days (**Figs. 4A–F, 6B–F**). Because nutrients are replenished over time in the human body, the continuous bioreactor was more suitable than the batch bioreactor for mimicking the nasal environment in our study [51]. A constant nutrient supply is crucial for bacterial growth, enabling them to adapt and respond to nutrients in the nasal environment [51]. After we inoculated batch and continuous bioreactors with the same nasal-swab specimens, *S. aureus* coexisted with other nasal bacteria (*e.g.*, *S. epidermidis* and *S. capitis*) in the continuous bioreactor (**Fig. 2B**), but not in the batch bioreactor (**Fig. 2A**). In the latter, *S. aureus* was the dominant species, comprising more than 97% of stable nasal microbiomes (**Fig. 2A**). This dominance eventually simplified the bacterial composition, posing limitations for nasal microbiome studies. We hypothesize, based on previous studies, that *S. aureus*’s dominance may be closely related to its high adaptability to the nutrient-limited environments of nasal habitats [52, 53]. *S. aureus* can fine-tune its expression of nutrient uptake systems to adapt to different environments [53]. Moreover, *S. aureus* has evolved to outcompete resident nasal bacteria by secreting antibacterial substances [52].

In a continuously fed bioreactor, other nasal bacteria can compete with *S. aureus* for nutrients, while antibacterial substances that were secreted into the medium are continuously washed out, resulting in lower concentrations than in a batch bioreactor. For example, *S. epidermis* established itself in our continuous bioreactor, even though a previous report noted that it did not grow well in the media that we used (SNM3) [14]. A likely reason is that co-colonization in the continuous bioreactor is favored, leading to the accumulation of growth factors from other community members that *S. epidermis* needs for optimal growth, and thereby preventing them from being washed out in our continuous-flow bioreactors.

Similarly, *in-vitro* culture models, which are regarded as batch systems, have been widely used to study nasal microbiomes to combat *S. aureus* infections [25, 53]. However, their closed system, which has restricted nutrients and lacks proper pH control, promotes *S. aureus* dominance and disrupts the maintenance of stable microbiomes [25, 53, 54]. Therefore, bacterial culture models are primarily used for short-term experiments lasting several days. In contrast, our continuous bioreactor, which was optimized to mimic the nasal environment, reliably cultivated nasal microbiomes for up to 36 days and prevented *S. aureus* dominance (**Figs. 4A–F, 6B–F**) [28]. We propose here the use of continuous bioreactors as a superior culture model for nasal microbiome studies.

### 4.2. Our continuous bioreactor is useful in devising *S. aureus* decolonization strategies from nasal microbiomes

We inoculated the bioreactor with nasal-swab specimens for optimization experiments **(Figs. 2A–E)**. However, nasal-swab specimens, which are *in-vivo* nasal microbiomes, make it challenging as an inoculum to predict the eventual microbial community structure of the stable nasal microbiomes. Because various microbes (*e.g.*, bacteria, archaea, fungi, and viruses) coexist in nasal-swab specimens, unpredictable polymicrobial interactions can affect the functioning of nasal microbiomes [3, 55]. Camus et al. and Navarro Diaz et al. addressed this problem by assembling SynComs comprising 5–10 nasal strains for an *in-vitro* culture model (*i.e.*, agar plates with SNM) [20, 21]. Isolated bacterial strains in the SynComs enable precise prediction and explanations of the composition and functioning of nasal microbiomes. To further investigate a possible *S. aureus* colonization strategy in the nasal microbiome, we inoculated a SynCom with 9 nasal strains into the optimized bioreactor, instead of using nasal-swab specimens (**Figs. 6A–F**).

Camus et al. found that tyrosine availability shapes *S. aureus* nasal colonization using SynComs [20]. The workers assessed *S. aureus* colonization using *in-vitro* cultures of nasal commensals and SynComs on SNM agar [20]. They identified that tyrosine auxotrophy plays an important role in the ability of *S. aureus* to develop within nasal communities. However, unstable nasal microbiomes (notably observed through *S. aureus* dominance in cultivated SynComs) and short experimental periods seem to limit the understanding of the biological mechanisms involved. Based on their findings, we employed our continuous bioreactor system to provide a complementary and more physiologically relevant approach to dissect the underlying mechanisms. Specifically, we cultivated two *S. aureus* strains with different abilities to metabolize tyrosine (Sa C or Sa A). We inoculated one of the two strains only after maintaining stable nasal microbiomes with SynCom C (**Fig. 6A**), which has low concentrations of amino acids (*i.e.*, tyrosine), and thereby imposing strong nutritional stress on *S. aureus* [20].

We found the tyrosine prototrophic strain Sa C to exhibit robust growth in the bioreactor with SynCom C (**Figs. 6C–D**). In contrast, Sa A, which is genetically auxotrophic to tyrosine, was rarely detected with SynCom C after inoculation (Days 20**–**36) (**Figs. 6E–F**). These results are consistent with the observations of Camus *et al.*, who demonstrated that SynCom C promotes Sa C proliferation while restricting that of Sa A during their co-culture experiments [20]. However, our system also highlights the community dynamics behind these effects, and particularly that Sa C colonization of this SynCom is associated with decreased abundances of *S. epidermidis* and almost complete elimination of *S. capitis*. This result yields two particularly noteworthy findings. First, using tyrosine auxotrophy may be a safe and effective approach for *S. aureus* decolonization in nasal microbiomes (**Figs. 6E–F**) [20], and should be further tested in our continuous bioreactor system. Second, to develop a nasal microbiome from SynComs that functions similarly to the original resident nasal microbiome, it is recommended to use isolated strains from a single volunteer. For example, SynCom C was able to proliferate only its own isolated strain (*i.e.*, Sa C). The effects and downstream applications of SynCom are most likely strain- and/or community-dependent (**Figs. 6B–F**).

With our continuous bioreactor, we can now consider various attempts to devise *S. aureus* decolonization strategies. For instance, we can inject various drugs and microbes into the bioreactors and evaluate their efficacy on *S. aureus* decolonization and microbiome changes throughout extended operating periods (**Figs. 6A–F**). We can also test various drug and microbial prescriptions by adjusting injection conditions (*e.g.*, start time, duration, and concentration). Furthermore, we can assess antimicrobial resistance mechanisms and the effects of drugs under more realistic conditions, which could inform the optimization of current treatments [56].

### 4.3. Our continuous bioreactor encountered numerous challenges

Our optimized bioreactor provides a physiologically relevant model for studying nasal microbiomes; however, we still face numerous challenges. One limitation is that we cannot cultivate all nasal bacteria in the bioreactor. The bacterial community compositions of inocula on Day 0 (*i.e.*, nasal-swab specimens and SynCom) and those of stable nasal microbiomes in the bioreactor were inconsistent (**Figs. 4A–F, 6B–F**). After inoculation with nasal-swab specimens into bioreactors, bacteria that had been below the detection limit in the inoculum (Day 0) proliferated during cultivation in the bioreactor (**Figs. 4C–D, S8C–D**). For instance, nasal-swab specimens from Volunteers 3 and 4, which were mainly composed of *Corynebacterium* spp., were found to be dominated by *Klebsiella* spp., *Bacillus* spp., and *Citrobacter* spp. at the end of the operating period (**Figs. 4C–D, S8C–D**). In addition, after inoculating the bioreactor with SynCom C, we did not detect nasal strains that required complete anaerobic conditions (*i.e.*, *Peptoniphilus harei* and *Anaerococcus octavius*) throughout the operating period (**Figs. 6B–F**). This is because selective pressures in the bioreactor environment may enrich certain nasal bacteria with outstanding adaptability to the bioreactor [31]. In addition, during bioreactor operation, the continuous flow of medium could wash away slow-growing bacteria before they adapt to the bioreactor environment [29]. Fortunately, the use of membranes and the recycling of some of these bacteria back into the bioreactor can be engineered.

The absence of human host cells in the current bioreactor can also create an environment that differs from that found in the human nose [57]. An organ-on-chip is a microfluidic device that enables the analysis of the structure, function, and physiology of living human organs [58]. Connecting the device to our bioreactor may be a solution, but few studies have been reported to date [24]. Moreover, we did not include innovative techniques to accurately measure and predict biological phenomena in the bioreactor. Deep functional analysis (*e.g.*, metatranscriptomics, metaproteomics, and multi-omics) can enhance our understanding of ecological niches and resistance mechanisms against *S. aureus* colonization in nasal microbiomes [1]. Mathematical and computational models are also necessary to systematically explain complex interactions in nasal microbiomes [59]. If these challenges are addressed in the future, our current bioreactor can serve as an improved model for studying *S. aureus* colonization of nasal microbiomes and for developing new strategies to limit it.

## 5. Conclusions

We developed a continuous bioreactor that is a more physiologically relevant model for studying nasal microbiomes than current *in-vitro* culture models. Our bioreactor setup, which was optimized by adjusting operating conditions (operating mode, dilution rate, temperature, pH, and medium composition), demonstrated high reproducibility across experiments and resilience to a pH perturbation. Furthermore, after inoculating our continuous bioreactor with six swab specimens and one SynCom, we developed stable nasal microbiomes for an extended operating period and assessed the ability of *S. aureus* strains to invade these communities. However, further efforts (*e.g.*, broad cultivation of nasal bacteria, integration of the human host, and in-depth functional analysis) are needed to develop a more physiologically relevant model for studying nasal microbiomes. Nevertheless, our findings suggest that our continuous bioreactor is a reasonable starting point for studying nasal microbiomes, which can complement the existing culture models.

## Supplementary information

**Additional file 1 (Supplementary figures): Fig. S1.** Standard curves of qPCR using 10-fold dilutions of genomic DNA. **A–B.** Standard curves for total bacteria (**A**) and *S. aureus* (**B**). E: efficiency. **Fig. S2.** Bacterial community compositions of six nasal-swab specimens from Volunteers 1–6. **A–B.** Relative abundances of the top ten genera (**A**) or species (**B**) in the nasal-swab specimens. The order of genera or species is based on the average relative abundance for the nasal-swab specimens from six volunteers. The most abundant genera or species are placed from bottom to top of the bar, but their names are listed from top to bottom in the legend. The remaining genera or species are grouped under ‘others’ when not in the top ten. **Fig. S3.** Optical densities of nasal microbiomes in the bioreactors throughout the operating period under various conditions for the optimization experiment. **A–E.** Operating mode (**A**), dilution rate (**B**), temperature (**C**), pH (**D**), and medium composition (**E**). SNM: synthetic nasal medium. **Fig. S4.** Bacterial community compositions of nasal-swab specimens and nasal microbiomes in the bioreactor under various conditions for the optimization experiment. **A–E.** Operating mode (**A**), dilution rate (**B**), temperature (**C**), pH (**D**), and medium composition (**E**). Nasal microbiomes were collected at the end of the operating period. The order of species for panels **A–C** is based on the average relative abundance of inoculum and nasal microbiomes in four bioreactors, which were operated in duplicate. The order of species for panels **D–E** is based on the average relative abundance of inoculum and nasal microbiomes in six bioreactors, which were operated in duplicate. For panels **A–E**, the most abundant species are placed from bottom to top of the bar, but their names are listed from top to bottom in the legend. However, the color for each species is the same between the panels. The remaining species not in the top ten are grouped under ‘others’. Ino.: inoculum, B: batch, C: continuous, SNM: synthetic nasal medium. **Fig. S5.** Copy numbers of *S. aureus* in nasal microbiomes in the bioreactor under various conditions for the optimization experiment. **A–E.** Operating mode (**A**), dilution rate (**B**), temperature (**C**), pH (**D**), and medium composition (**E**). Nasal microbiomes were collected at the end of the operating period. Error bars indicate standard deviations for three replicates. Statistical significance was determined by two-way ANOVA with Tukey’s multiple comparison test: *P<0.05, **P<0.01, ***P<0.001, ****P<0.0001. B: batch mode, C: continuous mode, SNM: synthetic nasal medium, P: adjusted p-value. **Fig. S6.** Reproducible comparison of bacterial community compositions between bioreactors inoculated with different nasal-swab specimens for the reproducibility experiment. **A–B.** Pearson correlation coefficient between the relative abundance of nasal-swab specimens 1 and 6 (**A**) and Nasal microbiomes 1 and 6 in the bioreactors at the end of the operating period (**B**). **C–D.** Relative abundances of the nasal microbiomes in the bioreactors that were inoculated with the nasal-swab specimen 1 (**C**) and 6 (**D**) throughout the operating period. Bacterial composition of the nasal-swab specimens is shown on Day 0 as an initial inoculum. The order and color of species for panels **C–D** are the same and are based on the combined average relative abundance throughout the operating period for both bioreactors. The most abundant species are placed from bottom to top of the bar, but their names are listed from top to bottom in the legend. The remaining species are grouped under ‘others’ when not in the top ten. Nas. Micr: Nasal microbiome. **Fig. S7.** Performance changes in the bioreactor with Nasal microbiomes A–F for the stability experiment. **A.** Optical densities. **B–C.** Copy numbers of total bacteria (**B**) and copy numbers of *S. aureus* (**C**) throughout the operating period. Nas. Micr.: Nasal microbiome. **Fig. S8.** Bacterial community compositions of Nasal microbiomes A–F in the bioreactor inoculated with six different nasal-swab specimens for the stability experiment. **A–F.** Relative abundances of the top ten genera in Nasal microbiome A (**A**), Nasal microbiome B (**B**), Nasal microbiome C (**C**), Nasal microbiome D (**D**), Nasal microbiome E (**E**), and Nasal microbiome F (**F**) throughout the operating period. Bacterial composition of the nasal-swab specimens is shown on Day 0 as an initial inoculum. The order of genera differs for each panel and is based on the combined average relative abundance throughout the operating period for each bioreactor. The most abundant genera are placed from bottom to top of the bar, but their names are listed from top to bottom in the legend. However, the color for each genus is the same between the panels. The remaining genera are grouped under ‘others’ when not in the top ten. **Fig. S9.** Nasal microbiome changes in the bioreactor inoculated with SynCom and different *S. aureus* strains for the decolonization experiment. SynCom C was inoculated into the bioreactor on Day 0, and two different *S. aureus* strains (Sa C or Sa A) were added on Day 20. **A.** Optical densities. **B–C.** Copy numbers of total bacteria (**B**) and copy numbers of *S. aureus* (**C**) throughout the operating period.

**Additional file 2 (Supplementary tables): Table S1.** Information on the six volunteers who participated in this study. **Table S2.** Information on the nasal strains used for SynCom C. **Table S3.** Schedule of the continuous bioreactor to cultivate nasal microbiomes through five experiments. **Table S4.** List of qPCR primers.

## Abbreviations

SynCom: Synthetic microbial community
SNM: Synthetic nasal medium
OD: Optical density
qPCR: Quantitative polymerase chain reaction
PCA: Principal coordinates analysis
MS: Mass spectrometry
AGC: Automatic gain control

## Declarations

### Ethics approval and consent to participate

This study was approved by the University of Tübingen (No. 109/2009 BO2), and informed consent was obtained from all volunteers.

### Consent for publication

Not applicable

### Availability of data and material

The datasets generated and analyzed during the current study are available in the NCBI BioProject repository under accession PRJNA1414135.

## Authors’ contributions

S.H. operated bioreactors and performed qPCR, and amplicon sequencing. M.N-D. and P.S. conducted non-targeted metabolomics. L.C. collected nasal-swab specimens from volunteers and provided SynComs. T.N.L. analyzed sequencing data. S.He., H.L., D.P., D.H.H., and L.T.A. designed the study. S.H. and L.T.A. wrote the manuscript. All authors read, reviewed, and approved the final manuscript.

## Acknowledgements

We acknowledge volunteers who participated in this study. We thank Nicolai Kreitli, Dr. Julia Schumacher, and Prof. Dr. Bastian Molitor at the University of Tübingen for their help in operating the bioreactors. We thank Jessica Franz at the University of Tübingen for building SynCom C. Special thanks to Dr. Libera Lo Presti at the University of Tübingen for reviewing the manuscript. Finally, we thank Dr. Sang-Hoon Lee and Prof. Hee-Deung Park at Korea University for providing comments about qPCR.

## References

1. Aggarwal N, Kitano S, Puah GRY, Kittelmann S, Hwang IY, Chang MW. Microbiome and human health: current understanding, engineering, and enabling technologies. Chem Rev. 2022;123(1):31–72. 10.1021/acs.chemrev.2c00431.

2. Yan M, Pamp SJ, Fukuyama J, Hwang PH, Cho D-Y, Holmes S, et al. Nasal microenvironments and interspecific interactions influence nasal microbiota complexity and *S. aureus* carriage. Cell Host Microbe. 2013;14(6):631–640. 10.1016/j.chom.2013.11.005.

3. Rawls M, Ellis AK. The microbiome of the nose. Annals of Allergy, Ann Allergy Asthma Immunol. 2019;122(1):17–24. 10.1016/j.anai.2018.05.009.

4. Belizário JE, Napolitano M. Human microbiomes and their roles in dysbiosis, common diseases, and novel therapeutic approaches. Front Microbiol. 2015;6:1050. 10.3389/fmicb.2015.01050.

5. Cole AL, Sundar M, Lopez A, Forsman A, Yooseph S, Cole AM. Identification of nasal Gammaproteobacteria with potent activity against *Staphylococcus aureus*: novel insights into the “noncarrier” state. mSphere. 2021;6(1):e01015–20. 10.1128/mSphere.01015-20.

6. Tai J, Han MS, Kwak J, Kim TH. Association between microbiota and nasal mucosal diseases in terms of immunity. Int J Mol Sci. 2021;22(9):4744. 10.3390/ijms22094744.

7. Hardy BL, Merrell DS. Friend or foe: interbacterial competition in the nasal cavity. J Bacteriol. 2021;203(5):e00480–20. 10.1128/JB.00480-20.

8. Aggarwal D, Bellis KL, Blane B, De Goffau MC, Wagner J, Ng DY, et al. Large-scale characterisation of the nasal microbiome redefines *Staphylococcus aureus* colonisation status. Nat Commun. 2025;16(1):10415. 10.1038/s41467-025-66564-4.

9. Zhang Y, Liu Z, Yuan F, Huang X, Wu D. Distinct inflammatory patterns and nasal bacterial dysbiosis in uncontrolled chronic rhinosinusitis. Eur Arch Otorhinolaryngol. 2025;282(6):3085–3096. 10.1007/s00405-025-09376-y.

10. Lee K, Pletcher SD, Lynch SV, Goldberg AN, Cope EK. Heterogeneity of microbiota dysbiosis in chronic rhinosinusitis: potential clinical implications and microbial community mechanisms contributing to sinonasal inflammation. Front Cell Infect Microbiol. 2018;8:168. 10.3389/fcimb.2018.00168.

11. Biswas K, Hoggard M, Jain R, Taylor MW, Douglas RG. The nasal microbiota in health and disease: variation within and between subjects. Front Microbiol. 2015;6:134. 10.3389/fmicb.2015.00134.

12. Wertheim HF, Melles DC, Vos MC, Van Leeuwen W, Van Belkum A, Verbrugh HA, et al. The role of nasal carriage in *Staphylococcus aureus* infections. Lancet Infect Dis. 2005;5(12):751–762. 10.1016/S1473-3099(05)70295-4.

13. Piewngam P, Otto M. *Staphylococcus aureus* colonisation and strategies for decolonisation. Lancet Microbe. 2024;5(6):e606–e618. 10.1016/S2666-5247(24)00040-5.

14. Krismer B, Liebeke M, Janek D, Nega M, Rautenberg M, Hornig G, et al. Nutrient limitation governs *Staphylococcus aureus* metabolism and niche adaptation in the human nose. PLoS Pathog. 2014;10(1):e1003862. 10.1371/journal.ppat.1003862.

15. Laux C, Peschel A, Krismer B. *Staphylococcus aureus* colonization of the human nose and interaction with other microbiome members. Microbiol Spectr. 2019;7:2. 10.1128/microbiolspec.GPP3-0029-2018.

16. Coates T, Bax R, Coates A. Nasal decolonization of *Staphylococcus aureus* with mupirocin: strengths, weaknesses and future prospects. J Antimicrob Chemother. 2009;64(1):9–15. 10.1093/jac/dkp159.

17. Baede VO, Barray A, Tavakol M, Lina G, Vos MC, Rasigade J-P, et al. Nasal microbiome disruption and recovery after mupirocin treatment in *Staphylococcus aureus* carriers and noncarriers. Sci Rep. 2022;12(1):19738. 10.1038/s41598-022-21453-4.

18. Edslev SM, Liu CM, Lo BZS, Meiniche H, Lilje B, Park DE, et al. Changes in nasal and throat microbiota composition during and after mupirocin and chlorhexidine decolonisation treatment in asymptomatic MRSA carriers: a longitudinal observational study. Clin Microbiol Infect. 2026. 10.1016/j.cmi.2026.02.020.

19. Vickery TW, Ramakrishnan VR, Suh JD. The role of *Staphylococcus aureus* in patients with chronic sinusitis and nasal polyposis. Curr Allergy Asthma Rep. 2019;19:21. 10.1007/s11882-019-0853-7.

20. Camus L, Franz J, Gerlach D, Lange A, Power JJ, Navarro-Díaz M, et al. Tyrosine availability shapes *Staphylococcus aureus* nasal colonization and interactions with commensal communities. bioRxiv. 2025. https://www.biorxiv.org/content/10.1101/2025.05.06.651429.

21. Navarro Diaz M, Camus L, Ham S, Angenent LT, Heilbronner S, Stincone P, et al. Community composition and strain identity drive metabolic competition and *Staphylococcus aureus* colonization resistance in synthetic nasal communities. bioRxiv 2026. https://www.biorxiv.org/content/10.64898/2026.01.08.698450.

22. Kiser KB, Cantey-Kiser JM, Lee JC. Development and characterization of a *Staphylococcus aureus* nasal colonization model in mice. Infect Immun. 1999, 67(10):5001–5006. 10.1128/IAI.67.10.5001-5006.1999.

23. Kokai-Kun JF. The cotton rat as a model for *Staphylococcus aureus* nasal colonization in humans: cotton rat *S. aureus* nasal colonization model. Methods Mol Biol. 2008;431:241–254. 10.1007/978-1-60327-032-8_19.

24. Biagini F, Daddi C, Calvigioni M, De Maria C, Zhang YS, Ghelardi E, et al. Designs and methodologies to recreate *in vitro* human gut microbiota models. Bio-Des Manuf. 2023;6:298–318. 10.1007/s42242-022-00210-6.

25. García Mendez DF, Sanabria J, Wist J, Holmes E. Effect of operational parameters on the cultivation of the gut microbiome in continuous bioreactors inoculated with feces: a systematic review. J Agric Food Chem. 2023;71(16):6213–6225. 10.1021/acs.jafc.2c08146.

26. Foo JL, Ling H, Lee YS, Chang MW. Microbiome engineering: current applications and its future. Biotechnol J. 2017;12(3):1600099. 10.1002/biot.201600099.

27. Hu H, Wang M, Huang Y, Xu Z, Xu P, Nie Y, et al. Guided by the principles of microbiome engineering: accomplishments and perspectives for environmental use. mLife. 2022;1(4):382–398. 10.1002/mlf2.12043.

28. Guzman-Rodriguez M, McDonald JA, Hyde R, Allen-Vercoe E, Claud EC, Sheth PM, et al. Using bioreactors to study the effects of drugs on the human microbiota. Methods. 2018;149:31–41. 10.1016/j.ymeth.2018.08.003.

29. Garcia Mendez DF, Egan S, Wist J, Holmes E, Sanabria J. Meta-analysis of the microbial diversity cultured in bioreactors simulating the gut microbiome. Microb Ecol. 2024;87:57. 10.1007/s00248-024-02369-0.

30. Auchtung JM, Robinson CD, Britton RA. Cultivation of stable, reproducible microbial communities from different fecal donors using minibioreactor arrays (MBRAs). Microbiome. 2015;3:42. 10.1186/s40168-015-0106-5.

31. Jin Z, Ng A, Maurice CF, Juncker D. The mini colon model: a benchtop multi-bioreactor system to investigate the gut microbiome. Gut Microbes. 2022;14(1):2096993. 10.1080/19490976.2022.2096993.

32. Krause JL, Schaepe SS, Fritz-Wallace K, Engelmann B, Rolle-Kampczyk U, Kleinsteuber S, et al. Following the community development of SIHUMIx-a new intestinal *in vitro* model for bioreactor use. Gut Microbes. 2020;11(4):1116–1129. 10.1080/19490976.2019.1702431.

33. Schäpe SS, Krause JL, Engelmann B, Fritz-Wallace K, Schattenberg F, Liu Z, et al. The simplified human intestinal microbiota (SIHUMIx) shows high structural and functional resistance against changing transit times in *in vitro* bioreactors. Microorganisms. 2019;7(12):641. 10.3390/microorganisms7120641.

34. Lawson CE, Harcombe WR, Hatzenpichler R, Lindemann SR, Löffler FE, O’Malley MA, et al. Common principles and best practices for engineering microbiomes. Nat Rev Microbiol. 2019;17:725–741. 10.1038/s41579-019-0255-9.

35. Lee S-H, Park J-H, Kang H-J, Lee YH, Lee TJ, Park H-D. Distribution and abundance of *Spirochaetes* in full-scale anaerobic digesters. Bioresour Technol. 2013;145:25–32. 10.1016/j.biortech.2013.02.070.

36. Anbazhagan D, Mui WS, Mansor M, Yan GOS, Yusof MY, Sekaran SD. Development of conventional and real-time multiplex PCR assays for the detection of nosocomial pathogens. Braz J Microbiol. 2011;42(2):448–458. 10.1590/S1517-83822011000200006.

37. Pfaffl MW. A new mathematical model for relative quantification in real-time RT-PCR. Nucleic Acids Res. 2001;29(9):e45. 10.1093/nar/29.9.e45.

38. Seol D, Lim JS, Sung S, Lee YH, Jeong M, Cho S, et al. Microbial identification using rRNA operon region: database and tool for metataxonomics with long-read sequence. Microbiol Spectr. 2022;10(2):e0201721. 10.1128/spectrum.02017-21.

39. Lucas TN, Biehain U, Gautam A, Gemeinhardt K, Lass T, Konzalla S, et al. MMonitor for real-time monitoring of microbial communities using long reads. Cell Rep Method. 2025;101266. 10.1016/j.crmeth.2025.101266.

40. Curry KD, Wang Q, Nute MG, Tyshaieva A, Reeves E, Soriano S, et al. Emu: species-level microbial community profiling of full-length 16S rRNA oxford nanopore sequencing data. Nat Methods. 2022;19:845–853. 10.1038/s41592-022-01520-4.

41. Stincone P, Pakkir Shah AK, Schmid R, Graves LG, Lambidis SP, Torres RR, et al. Evaluation of data-dependent MS/MS acquisition parameters for non-targeted metabolomics and molecular networking of environmental samples: focus on the Q exactive platform. Anal Chem. 2023;95(34):12673–12682. 10.1021/acs.analchem.3c01202.

42. Pakkir Shah AK, Walter A, Ottosson F, Russo F, Navarro-Diaz M, Boldt J, et al. Statistical analysis of feature-based molecular networking results from non-targeted metabolomics data. Nat Protoc. 2025;20(1):92–162. 10.1038/s41596-024-01046-3.

43. Gu Z, Eils R, Schlesner M. Complex heatmaps reveal patterns and correlations in multidimensional genomic data. Bioinfor. 2016;32(18):2847–2849. 10.1093/bioinformatics/btw313.

44. Liu CM, Price LB, Hungate BA, Abraham AG, Larsen LA, Christensen K, et al. *Staphylococcus aureus* and the ecology of the nasal microbiome. Sci Adv. 2015;1(5):e1400216. 10.1126/sciadv.1400216.

45. Hehar S, Mason J, Stephen A, Washington N, Jones N, Jackson S, et al. Twenty four hour ambulatory nasal pH monitoring. Clin Otolaryngol Allied Sci. 1999;24(1):24–25. 10.1046/j.1365-2273.1999.00190.x.

46. Washington N, Steele R, Jackson S, Bush D, Mason J, Gill D, et al. Determination of baseline human nasal pH and the effect of intranasally administered buffers. Int J Pharm. 2000;198(2):139–146. 10.1016/s0378-5173(99)00442-1.

47. Dedrick S, Akbari MJ, Dyckman SK, Zhao N, Liu Y-Y, Momeni B. Impact of temporal pH fluctuations on the coexistence of nasal bacteria in an *in silico* community. Front Microbiol. 2021;12:613109. 10.3389/fmicb.2021.613109.

48. Mougi A. pH adaptation stabilizes bacterial communities. NPJ Biodivers. 2024;3(1):32. 10.1038/s44185-024-00063-5.

49. Neugent ML, Hulyalkar NV, Nguyen VH, Zimmern PE, De Nisco NJ. Advances in understanding the human urinary microbiome and its potential role in urinary tract infection. MBio. 2020;11(2): e00218–20. 10.1128/mBio.00218-20.

50. Keck T, Leiacker R, Riechelmann H, Rettinger G. Temperature profile in the nasal cavity. Laryngoscope. 2000;110(4):651–654. 10.1097/00005537-200004000-00021.

51. Brown SA, Palmer KL, Whiteley M. Revisiting the host as a growth medium. Nat Rev Microbiol. 2008;6(9):657–666. 10.1038/nrmicro1955.

52. Garrett SR, Palmer T. The role of proteinaceous toxins secreted by *Staphylococcus aureus* in interbacterial competition. FEMS Microbes. 2024;5:xtae006. 10.1093/femsmc/xtae006.

53. Krismer B, Weidenmaier C, Zipperer A, Peschel A. The commensal lifestyle of *Staphylococcus aureus* and its interactions with the nasal microbiota. Nat Rev Microbiol. 2017;15(11):675–687. 10.1038/nrmicro.2017.104.

54. Kumar S, Wittmann C, Heinzle E. Minibioreactors. Biotechnol Lett. 2004;26(1):1–10. 10.1023/b:bile.0000009469.69116.03.

55. Peters BM, Jabra-Rizk MA, O’May GA, Costerton JW, Shirtliff ME. Polymicrobial interactions: impact on pathogenesis and human disease. Clin Microbiol Rev. 2012;25(1):193–213. 10.1128/CMR.00013-11.

56. Udekwu KI, Levin BR. *Staphylococcus aureus* in continuous culture: a tool for the rational design of antibiotic treatment protocols. PLoS One. 2012;7(7):e38866. 10.1371/journal.pone.0038866.

57. Huang S, Hon K, Bennett C, Hu H, Menberu M, Wormald P-J, et al. *Corynebacterium accolens* inhibits *Staphylococcus aureus* induced mucosal barrier disruption. Front Microbiol. 2022;13:984741. 10.3389/fmicb.2022.984741.

58. Bein A, Shin W, Jalili-Firoozinezhad S, Park MH, Sontheimer-Phelps A, Tovaglieri A, et al. Microfluidic organ-on-a-chip models of human intestine. Cell Mol Gastroenterol Hepatol. 2018;5(4):659–668. 10.1016/j.jcmgh.2017.12.010.

59. Deek RA, Ma S, Lewis J, Li H. Statistical and computational methods for integrating microbiome, host genomics, and metabolomics data. eLife. 2024;13:e88956. 10.7554/eLife.88956.

