## Supplementary material for "Development of a continuous bioreactor to maintain stable nasal microbiomes from swab specimens and synthetic communities": Fig. S

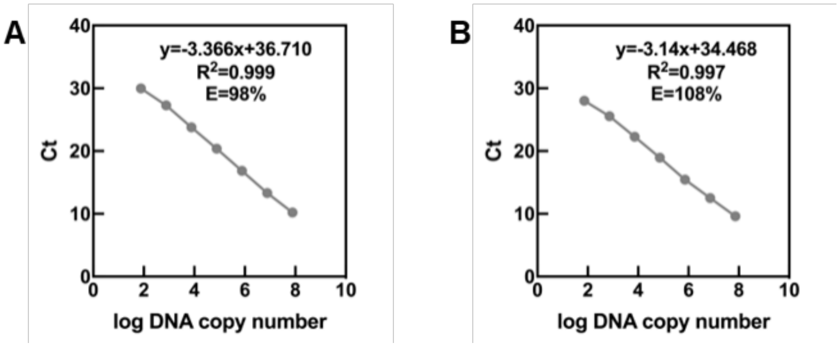


**Fig. S1.** Standard curves of qPCR using 10-fold dilutions of genomic DNA. **A–B.** Standard curves for total bacteria (**A**) and *S. aureus* (**B**). E: efficiency.

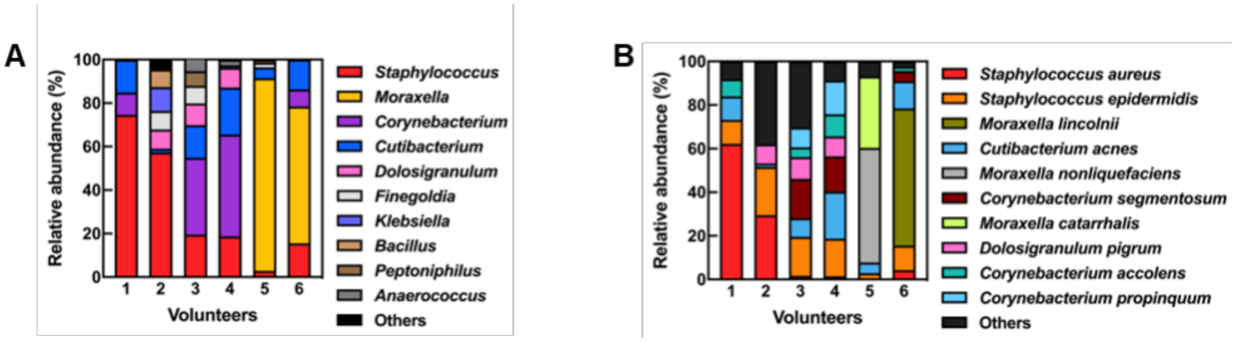


**Fig. S2.** Bacterial community compositions of six nasal-swab specimens from Volunteers 1–6. **A–B.** Relative abundances of the top ten genera (**A**) or species (**B**) in the nasal-swab specimens. The order of genera or species is based on the average relative abundance for the nasal-swab specimens from six volunteers. The most abundant genera or species are placed from bottom to top of the bar, but their names are listed from top to bottom in the legend. The remaining genera or species are grouped under 'others' when not in the top ten.

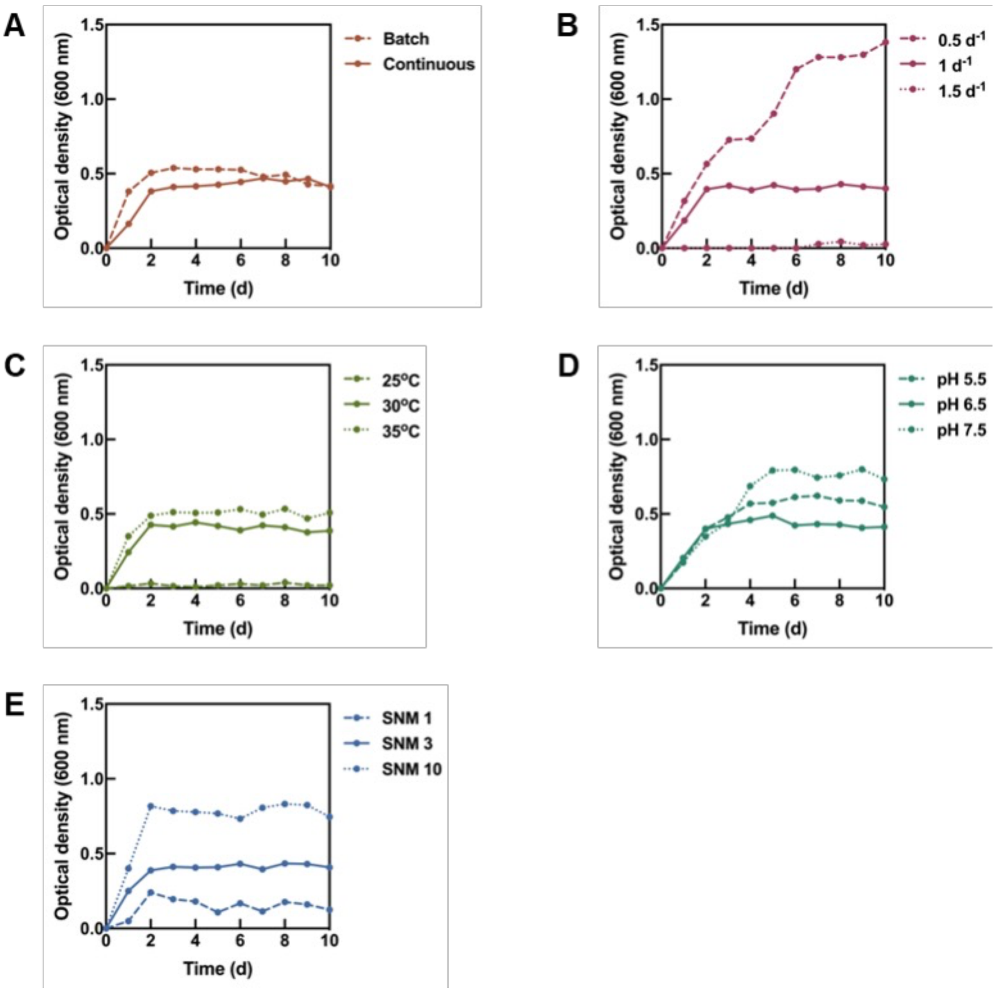


**Fig. S3.** Optical densities of nasal microbiomes in the bioreactors throughout the operating period under various conditions for the optimization experiment. **A–E.** Operating mode (**A**), dilution rate (**B**), temperature (**C**), pH (**D**), and medium composition (**E**). SNM: synthetic nasal medium.

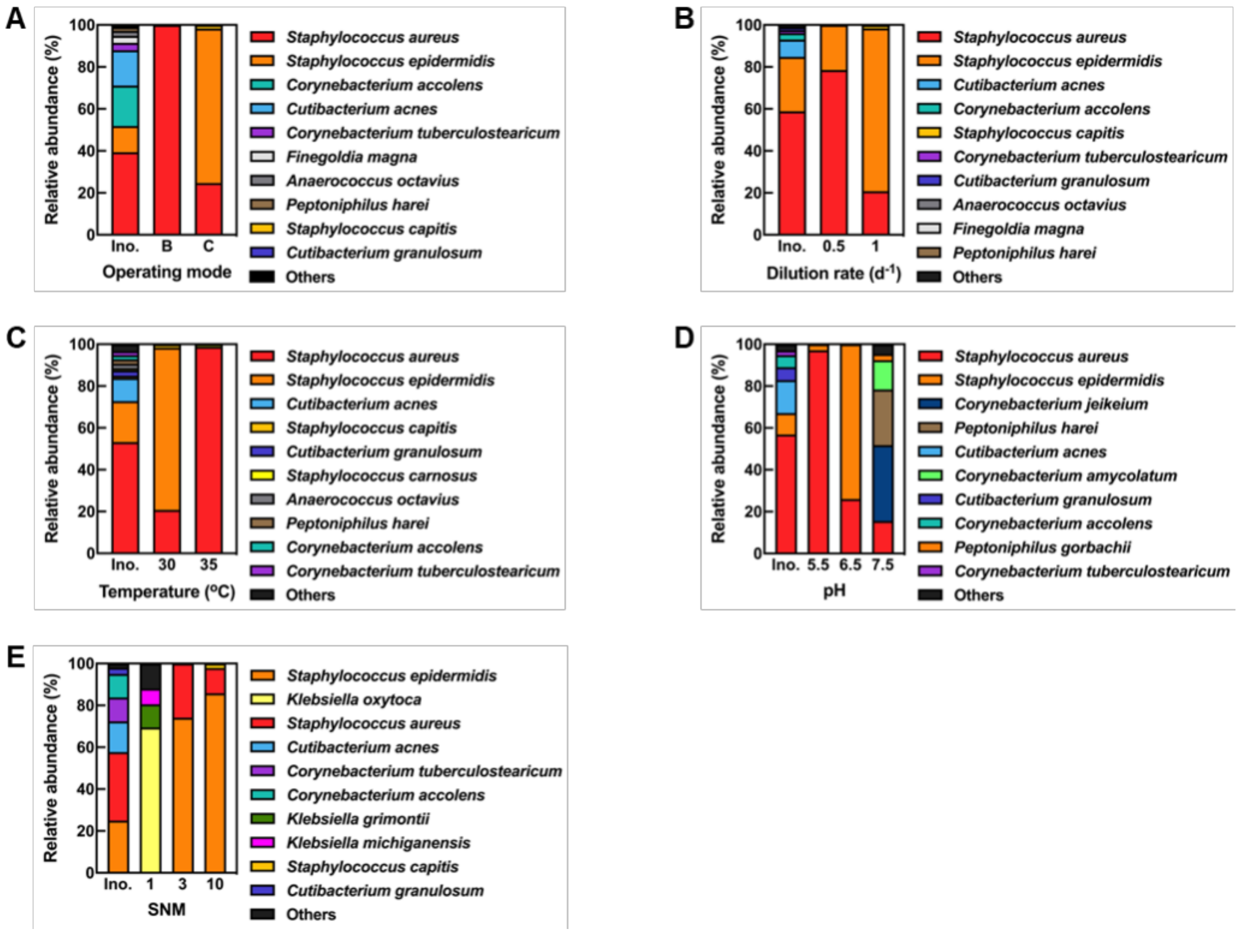


**Fig. S4.** Bacterial community compositions of nasal-swab specimens and nasal microbiomes in the bioreactor under various conditions for the optimization experiment. **A–E.** Operating mode (**A**), dilution rate (**B**), temperature (**C**), pH (**D**), and medium composition (**E**). Nasal microbiomes were collected at the end of the operating period. The order of species for panels **A–C** is based on the average relative abundance of inoculum and nasal microbiomes in four bioreactors, which were operated in duplicate. The order of species for panels **D–E** is based on the average relative abundance of inoculum and nasal microbiomes in six bioreactors, which were operated in duplicate. For panels **A–E**, the most abundant species are placed from bottom to top of the bar, but their names are listed from top to bottom in the legend. However, the color for each species is the same between the panels. The remaining species not in the top ten are grouped under 'others'. Ino.: inoculum, B: batch, C: continuous, SNM: synthetic nasal medium.

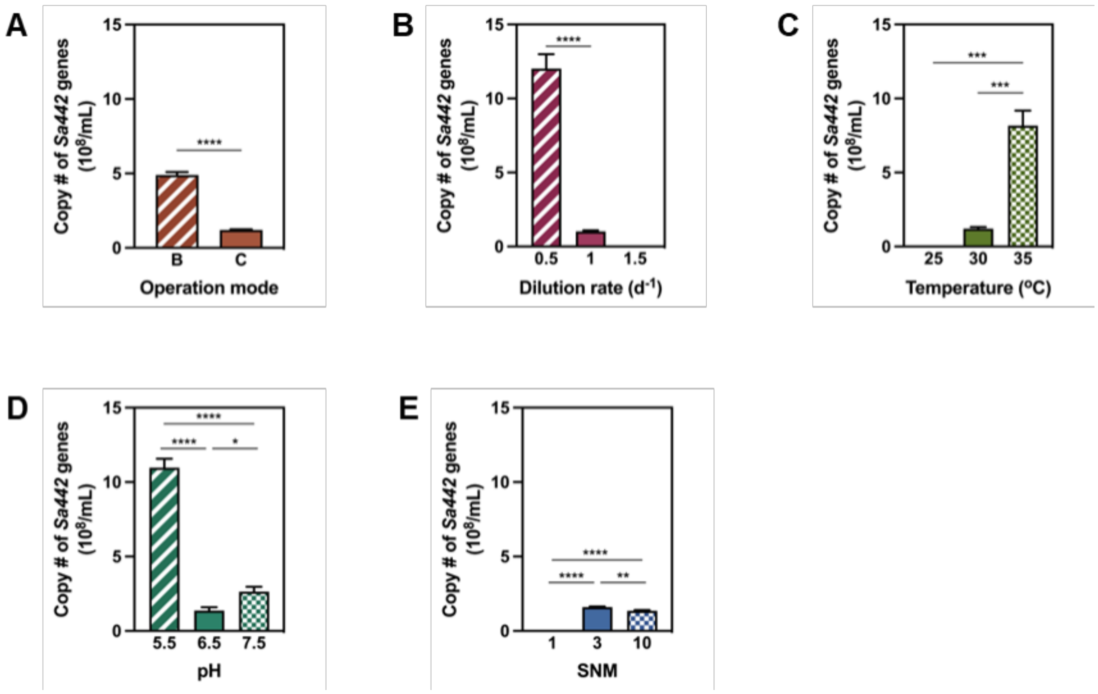


**Fig. S5.** Copy numbers of *S. aureus* in nasal microbiomes in the bioreactor under various conditions for the optimization experiment. **A–E.** Operating mode (**A**), dilution rate (**B**), temperature (**C**), pH (**D**), and medium composition (**E**). Nasal microbiomes were collected at the end of the operating period. Error bars indicate standard deviations for three replicates. Statistical significance was determined by two-way ANOVA with Tukey's multiple comparison test: $*P<0.05$, $**P<0.01$, $***P<0.001$, $****P<0.0001$. B: batch mode, C: continuous mode, SNM: synthetic nasal medium, P: adjusted p-value.

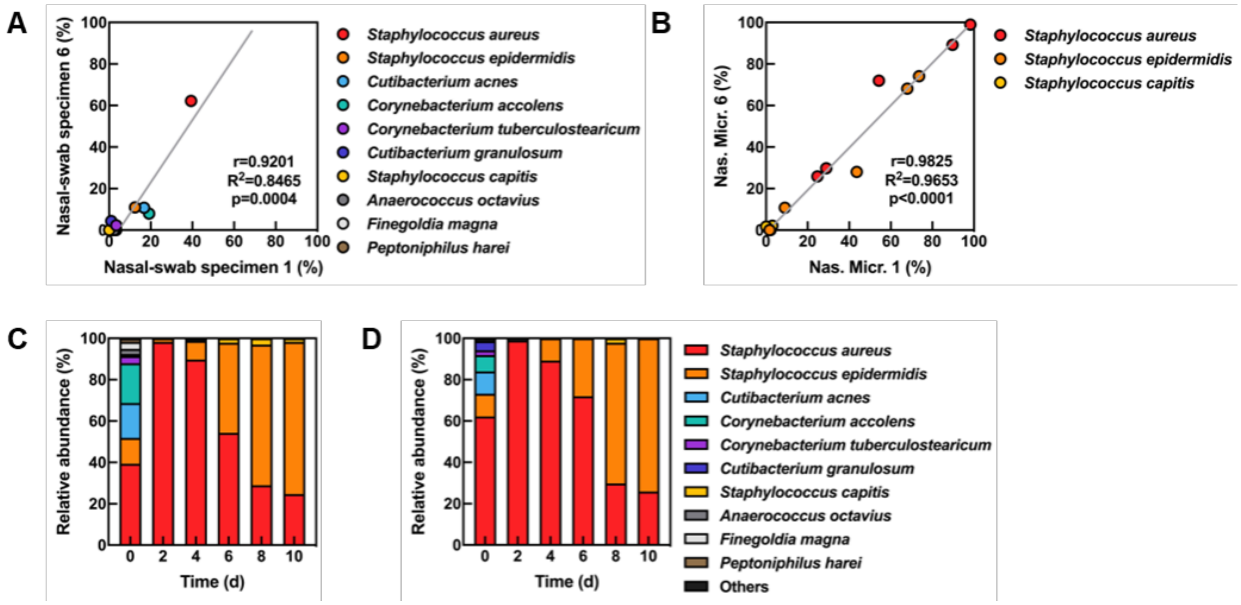


**Fig. S6.** Reproducible comparison of bacterial community compositions between bioreactors inoculated with different nasal-swab specimens for the reproducibility experiment. **A–B.** Pearson correlation coefficient between the relative abundance of nasal-swab specimens 1 *and* 6 (**A**) and Nasal microbiomes 1 *and* 6 in the bioreactors at the end of the operating period (**B**). **C–D.** Relative abundances of the nasal microbiomes in the bioreactors that were inoculated with the nasal-swab specimen 1 (**C**) *and* 6 (**D**) throughout the operating period. Bacterial composition of the nasal-swab specimens is shown on Day 0 as an initial inoculum. The order and color of species for panels **C–D** are the same and are based on the combined average relative abundance throughout the operating period for both bioreactors. The most abundant species are placed from bottom to top of the bar, but their names are listed from top to bottom in the legend. The remaining species are grouped under 'others' when not in the top ten. Nas. Micr: Nasal microbiome.

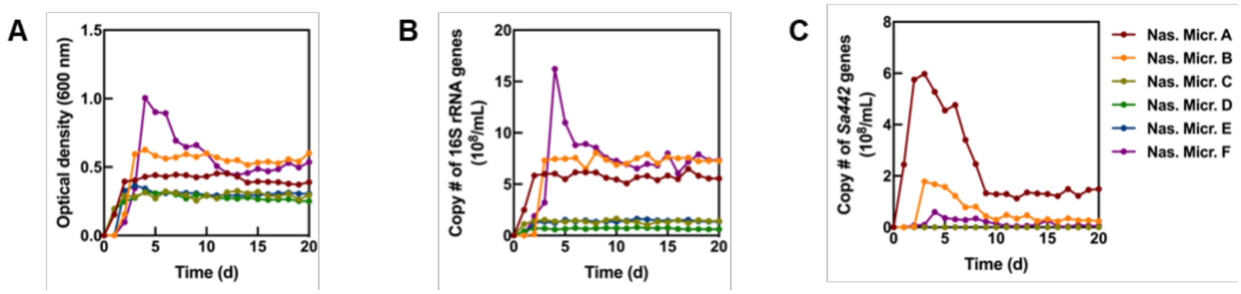


**Fig. S7.** Performance changes in the bioreactor with Nasal microbiomes A–F for the stability experiment. **A.** Optical densities. **B–C.** Copy numbers of total bacteria (**B**) and copy numbers of *S. aureus* (**C**) throughout the operating period. Nas. Micr.: Nasal microbiome.

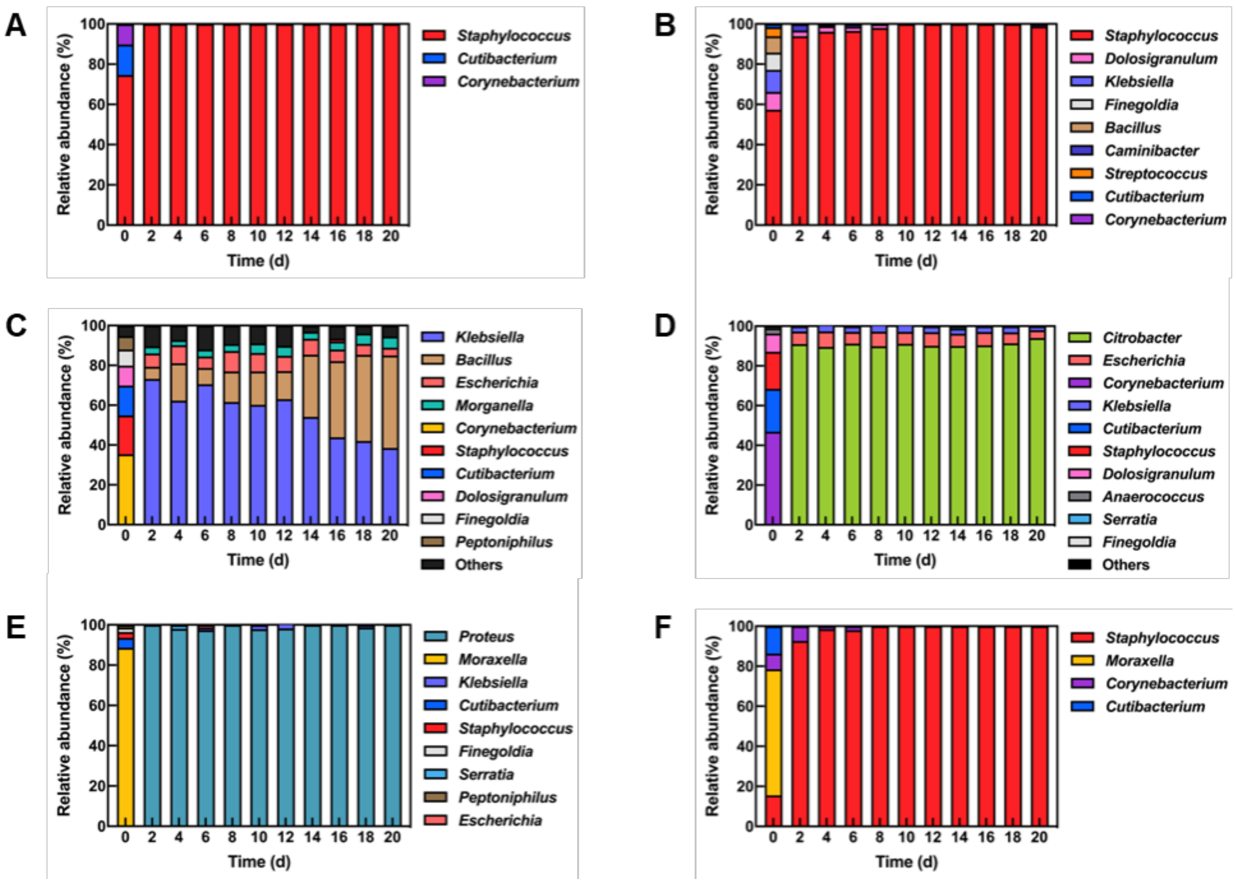


**Fig. S8.** Bacterial community compositions of Nasal microbiomes A–F in the bioreactor inoculated with six different nasal-swab specimens for the stability experiment. **A–F.** Relative abundances of the top ten genera in Nasal microbiome A (**A**), Nasal microbiome B (**B**), Nasal microbiome C (**C**), Nasal microbiome D (**D**), Nasal microbiome E (**E**), and Nasal microbiome F (**F**) throughout the operating period. Bacterial composition of the nasal-swab specimens is shown on Day 0 as an initial inoculum. The order of genera differs for each panel and is based on the combined average relative abundance throughout the operating period for each bioreactor. The most abundant genera are placed from bottom to top of the bar, but their names are listed from top to bottom in the legend. However, the color for each genus is the same between the panels. The remaining genera are grouped under ‘others’ when not in the top ten.

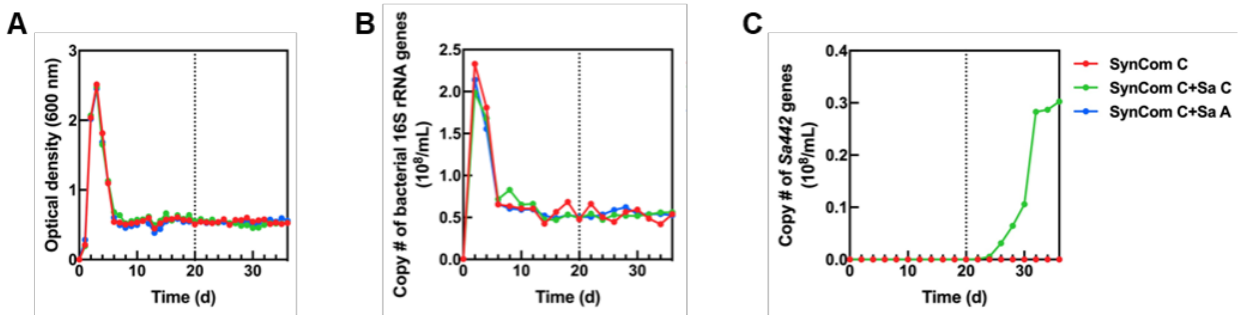


**Fig. S9.** Nasal microbiome changes in the bioreactor inoculated with SynCom and different *S. aureus* strains for the decolonization experiment. SynCom C was inoculated into the bioreactor on Day 0, and two different *S. aureus* strains (Sa C or Sa A) were added on Day 20. **A.** Optical densities. **B–C.** Copy numbers of total bacteria (**B**) and copy numbers of *S. aureus* (**C**) throughout the operating period.
