## Supplementary material for "Development of a continuous bioreactor to maintain stable nasal microbiomes from swab specimens and synthetic communities": Table S

**Table S1.** Information on the six volunteers who participated in this study.

| Vol. | General characteristics | | Sequencing | |
|---|---|---|---|---|
| | Gender | Age | Most abundant genera | Relative *S. aureus* abundance (%) |
| 1 | Female | 30–40 | *Staphylococcus* spp. | 62 |
| 2 | Female | 20–30 | *Staphylococcus* spp. | 29 |
| 3 | Male | 20–30 | *Corynebacterium* spp. | 2 |
| 4 | Male | 20–30 | *Corynebacterium* spp. | 1 |
| 5 | Male | 40–50 | *Moraxella* spp. | 0 |
| 6 | Female | 40–50 | *Moraxella* spp. | 4 |

**Table S2.** Information on the nasal strains used for SynCom C.

| Species | LaCa collection[A] | | Culture conditions | | | SynCom |
|---|---|---|---|---|---|---|
| | Identifier | Source[B] | Medium[C] | Duration | Oxygenation | |
| *Staphylococcus aureus* | Sa A | Vol. A | BHI | Overnight | Aerobiosis | Excluded |
| *Staphylococcus aureus* | Sa C | Vol. C | BHI | Overnight | Aerobiosis | Excluded |
| *Staphylococcus epidermidis* | Se C | Vol. C | BHI | Overnight | Aerobiosis | Included |
| *Staphylococcus capitis* | Sc C | Vol. C | BHI | Overnight | Aerobiosis | Included |
| *Micrococcus luteus* | Ml C | Vol. C | BHI | Overnight | Aerobiosis | Included |
| *Corynebacterium tuberculostearicum* | Ct C | Vol. C | BHI + 0.2% Tween 80 | 1 day | Aerobiosis | Included |
| *Cutibacterium avidum* | Cv C | Vol. C | TSB | 2 days | Anaerobiosis | Included |
| *Cutibacterium acnes* | Cc C | Vol. C | TSB | 4 days | Anaerobiosis | Included |
| *Cutibacterium granulosum* | Cg C | Vol. C | TSB | 4 days | Anaerobiosis | Included |
| *Anaerococcus octavius* | Ao C | Vol. C | TSA + 5% sheep blood | 7 days | Anaerobiosis | Included |
| *Peptoniphilus harei* | Ph C | Vol. C | TSA + 5% sheep blood | 7 days | Anaerobiosis | Included |

**[A]** LaCa collection was established by Camus et al. **[B]** Vol. C corresponds to Vol. 1 from this study. Vol.: volunteer. **[C]** BHI: brain heart infusion, SynCom: synthetic microbial community, TSB: tryptic soy broth, TSA: tryptic soy agar.

**Table S3.** Schedule of the continuous bioreactor to cultivate nasal microbiomes through five experiments.

| | | Time (d) | | | | | | | | | | | | | | | | | | |
|---|---|---|---|---|---|---|---|---|---|---|---|---|---|---|---|---|---|---|---|---|
| | | 0 | 2 | 4 | 6 | 8 | 10 | 12 | 14 | 16 | 18 | 20 | 22 | 24 | 26 | 28 | 30 | 32 | 34 | 36 |
| 1. Optimization experiment | 1-1. Reactor operation | | | | | | | | | | | | | | | | | | | |
| | 1-2. Inoculation | | | | | | | | | | | | | | | | | | | |
| | 1-3. OD, qPCR | | | | | | | | | | | | | | | | | | | |
| | 1-4. Sequencing | | | | | | | | | | | | | | | | | | | |
| 2. Reproducibility, perturbation, and stability experiments | 2-1. Reactor operation | | | | | | | | | | | | | | | | | | | |
| | 2-2. Inoculation | | | | | | | | | | | | | | | | | | | |
| | 2-3. OD, qPCR | | | | | | | | | | | | | | | | | | | |
| | 2-4. Sequencing | | | | | | | | | | | | | | | | | | | |
| | 2-5. Metabolomics | | | | | | | | | | | | | | | | | | | |
| 3. Decolonization experiment | 3-1. Reactor operation | | | | | | | | | | | | | | | | | | | |
| | 3-2. Inoculation | | | | | | | | | | | | | | | | | | | |
| | 3-3. OD, qPCR | | | | | | | | | | | | | | | | | | | |
| | 3-4. Sequencing | | | | | | | | | | | | | | | | | | | |

OD: optical density, qPCR: quantitative polymerase chain reaction

**Table S4.** List of qPCR primers.

| Primer | Target | Sequence | GC (%) | Tm (°C) |
|---|---|---|---|---|
| 27-F | Total bacteria | AGA GTT TGA TCC TGG CTC AG | 50 | 61 |
| 342-R | | CTG CTG CSY CCC GTA G | 63 | 58 |
| *Sa442*-F | *S. aureus* | TCG GTA CAC GAT ATT CTT CAC | 43 | 59 |
| *Sa442*-R | | ACT CTC GTA TGA CCA GCT TC | 50 | 59 |
